# Dissociating the contributions of sensorimotor striatum to automatic and visually-guided motor sequences

**DOI:** 10.1101/2022.06.13.495989

**Authors:** Kevin G. C. Mizes, Jack Lindsey, G. Sean Escola, Bence P. Ölveczky

## Abstract

The ability to sequence movements in response to new task demands enables rich and adaptive behavior. Such flexibility, however, is computationally costly and can result in halting performances. Practicing the same motor sequence repeatedly can render its execution precise, fast, and effortless, i.e., ‘automatic’. The basal ganglia are thought to underlie both modes of sequence execution, yet whether and how their contributions differ is unclear. We parse this in rats trained to perform the same motor sequence in response to cues and in an overtrained, or ‘automatic’, condition. Neural recordings in the sensorimotor striatum revealed a kinematic code independent of execution mode. While lesions affected the detailed kinematics similarly across modes, they disrupted high-level sequence structure for automatic, but not visually-guided, behaviors. These results suggest that the basal ganglia contribute to learned movement kinematics and are essential for ‘automatic’ motor skills but can be dispensable for sensory-guided motor sequences.

## Introduction

Our brain’s capacity to organize movements and actions in response to new challenges allows us to imitate trendy dance moves or play Chopin etudes from sheet music. Assembling motor sequences in such controlled and deliberate ways can be mentally taxing and computationally costly, resulting in slow^1^ and error-prone performances subject to cognitive interference^2,3^. However, executing the same motor sequence, such as typing a password or playing a favorite piano sonata, repeatedly and consistently, can turn it into a continuous task-specific movement pattern that is fluid, fast^4–7^, precise^5^, efficient^8^, and less cognitively demanding^2,9,10^; in a word: ‘automatic’^11–13^. Thus, the very same motor sequence can be executed in modes that are both qualitatively and subjectively distinct^1,12–14^.

Given that the specification of the same motor sequence can differ so markedly (Fig. 1), the underlying neural circuits, or the functions they implement, are thought to differ as well^10,15–21^. For example, a discrete motor sequence whose progression is informed by external sensory cues will engage a serial action selection process that likely engages higher-level circuits^22,23^ (Fig. 1a). Highly overtrained, or ‘automatic’, motor sequences, on the other hand, can be specified in terms of sequential low-level motor commands (Fig. 1c)^24–26^. Indeed, our colloquial reference to automatic behaviors being stored in ‘muscle memory’ reflects a subjective sense that they are, in contrast to sensory-guided motor sequences, less reliant on cognitive processes and produced by circuits closer to the motor periphery^11,14,15,19,27^.

**Figure 1:**
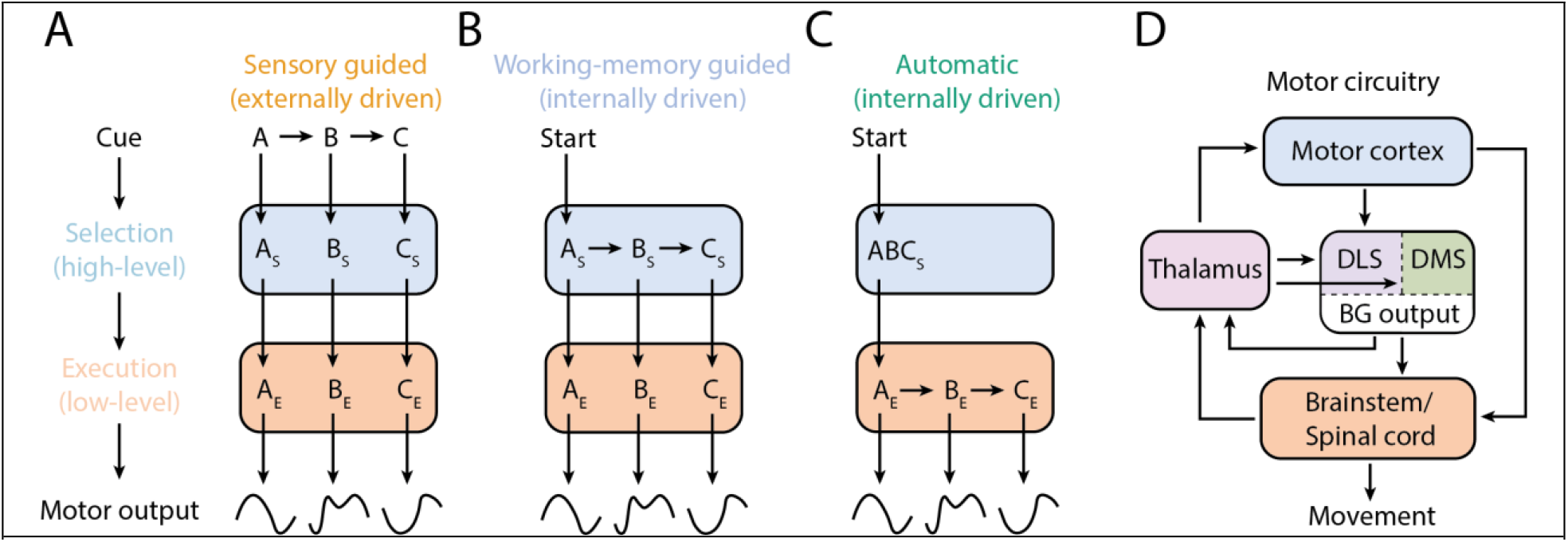
Conceptual schematic showing differences in how a motor sequence can be specified and produced. **A-C**. Motor control is thought to be hierarchical^32,40,41^. To perform a new discrete motor sequence (A-B-C), high-level circuits select and order the requisite motor elements (i.e., select A, then B, then C), while low-level circuits implement the detailed control of the movements. **A**. Sensory-guided and **B**. working memory-guided sequences likely engage higher-order selection-level processes, whereas **C**. automatic sequences can become consolidated and fully specified in low-level execution circuits^11,14,32^. Schematic adapted from^16,32^. **D**. Simplified schematic of motor circuits discussed in this study. The BG is the nexus of several intersecting motor pathways, allowing them to affect motor implementation by modulating both brainstem and motor cortical dynamics. Omitted, for clarity, are somatosensory and prefrontal projections, cerebellar inputs, and dopaminergic midbrain projections.

Besides being sensory-guided and automatic, motor sequences can also be informed by working memory (Fig. 1b), as is the case when we try to imitate our piano teacher or reproduce our own improvisations from a few moments ago. The generation of such motor sequences are akin to sensory-guided ones in that they too demand considerable mental effort^28^ and are informed by (remembered) sensory experiences. Given these shared qualities, sensory-guided and working memory guided sequences are often collectively referred to as ‘controlled’^1,13,16,17,29^ to distinguish them from automatic behaviors. However, working memory-guided motor sequences are also similar to automatic ones in that their progression is informed by internal neural processes rather than immediately available sensory cues. Hence these two modes of sequence execution, automatic and working memory-guided, are sometimes referred to as ‘internally’ generated, to distinguish them from ‘externally’ cued ones^20,22,30^.

Yet the degree to which the distinctions between ‘controlled’ vs ‘automatic’ and ‘externally vs ‘internally’ generated motor sequences map onto specific neural circuits and mechanisms is unclear^16,22,31,32^. Here we set out to probe how the neural implementations of automatic, visually- and working memory-guided sequences differ, focusing on the sensorimotor and associative arms of the basal ganglia (BG). While these pathways have been implicated in various aspects of motor sequence learning and execution^18,27,33–36^, their specific contribution to sensory-guided, working memory-guided, and automatic behaviors, have yet to be fully understood^15,19,37–39^.

Because sensory-guided motor sequences can be equated to serial action-selection or decision making^23,32,42^, it follows from BG’s acknowledged role in these processes^18,43–45^ that they should be central to generating such behaviors. While recordings showing sequence-specific neural activity in the primate and human BG are consistent with this view^46,47^, some recent studies have called this into question. For example, inactivating a main output nucleus of the BG (Globus Pallidus internal segment, GPi) in monkeys did not affect their ability to generate visually-guided reaching sequences beyond slightly reducing the vigor of the constituent movements^48^.

BG has also been implicated in working memory-guided sequences. Indeed, recordings from GPi neurons in monkeys performing the same sequences guided by visual cues and working memory revealed more task-modulated neurons in working memory trials^49,50^, an indication that the BG contribute differently to the two execution modes, a notion supported also by studies in humans^51^.

In contrast to sensory- and working memory-guided motor sequences, ‘automatic’ motor sequences lack any flexibility – once initiated, the progression of the behavior is fixed, obviating the need for discrete action selection. Studies showing neural activity in sensorimotor regions of the BG preferentially encoding the start and stop of overtrained behaviors^52–56^ imply that repeated practice transforms an initially controlled motor sequence – with distinct elements and choice points – into an immutable continuous sequential movement pattern^1,12^. While sensorimotor regions of the BG are thought to help select and initiate such ‘chunked’ behaviors^57–59^, the implication is that their detailed progression and specification are elaborated in downstream circuits.

The generality of this view, however, was challenged by recent studies showing that the BG is necessary for generating the detailed kinematics of stereotyped learned behaviors^24,60–63^. In these studies, activity patterns in the sensorimotor striatum did not reflect the starts and stops of the overtrained behaviors, nor specific choice points in the sequences; rather, the neurons encoded the kinematic details of the sequential behaviors, suggesting an essential role for the BG in the continuous low-level kinematic specification of automatic behaviors (Fig 1c).

To get at the distinction in how the BG contribute to the different types of motor sequences, we designed a task for rats that trains them to perform the very same motor sequence in the three execution modes discussed above (Fig 1). In the first mode, each element in the sequence is cued by a visual stimulus (‘sensory-guided’); in the second, the sequence is informed by working memory (‘working memory-guided); and in the third, the sequence is ‘automatic’ thanks to lengthy overtraining on the same sequence. Using the pianist as an analogy, the first is akin to playing a piece from sheet music, the second is repeating that very piece from memory, and the third is the condition that emerges after practicing it diligently for many weeks.

To distinguish the contributions of the BG to these different execution modes, we focused on the dorsal striatum, the input zone to the sensorimotor and associative arms of the BG. Surprisingly, neural recordings in sensorimotor striatum (dorsolateral striatum, DLS, in rodents) – a region implicated in behavioral automaticity^11,64^ – revealed no meaningful difference across the execution modes, with neurons representing low-level kinematic features in all cases. The neural population showed no selectivity for higher-level attributes of the behavior, such as the order or sequential context of a given movement.

Lesions to the DLS, however, revealed a stark contrast across execution modes, with sequence organization being essentially lost for automatic and working memory-guided sequences, but largely preserved for sequences informed by visual cues. Additionally, the movements across all three conditions were slower and more variable post-lesion, resembling the animal’s engagement with the task early in learning, a finding consistent with a general role for the BG in specifying the detailed kinematics, including the vigor^24,48^, of learned movements. Lesions of the associative region of the striatum (dorsomedial striatum in rodent; DMS), widely implicated in behavioral flexibility^65–69^, had no lasting effect on either the performance or the kinematics in either execution mode.

A network model simulating DLS and its interactions with other motor circuits recapitulated these results, including the similarity of DLS activity across execution modes and the differing impacts of DLS lesions on visually-guided and automatic sequences. The model also explained the shared role for DLS in the kinematic specification across execution modes by showing that differing inputs to DLS are transformed into common activity patterns that sculpt movement features through interactions with downstream control circuits.

## Results

### A discrete sequence production task for rats

To directly compare the neural substrates of visually-guided, working memory-guided, and automatic motor sequences, we designed, based on similar paradigms in humans and non-human primates (NHPs)^20,23,70^, a discrete sequence production task in which rats execute the same motor sequence in the three different modes.

To facilitate comparisons to other motor-related studies in rodents, including our own^24,25^, which probe forelimb^33,36,54,56,62,72^ and whole-body orienting^73–75^ movements, we opted for a ‘piano-playing’ task, in which rats are rewarded for performing sequences of three keypresses on a three-key ‘piano’ in a prescribed order, alternating between forelimb lever-presses and orienting movements (12 possible sequences; see Fig. 2a, Supplementary Movie 1).

**Figure 2:**
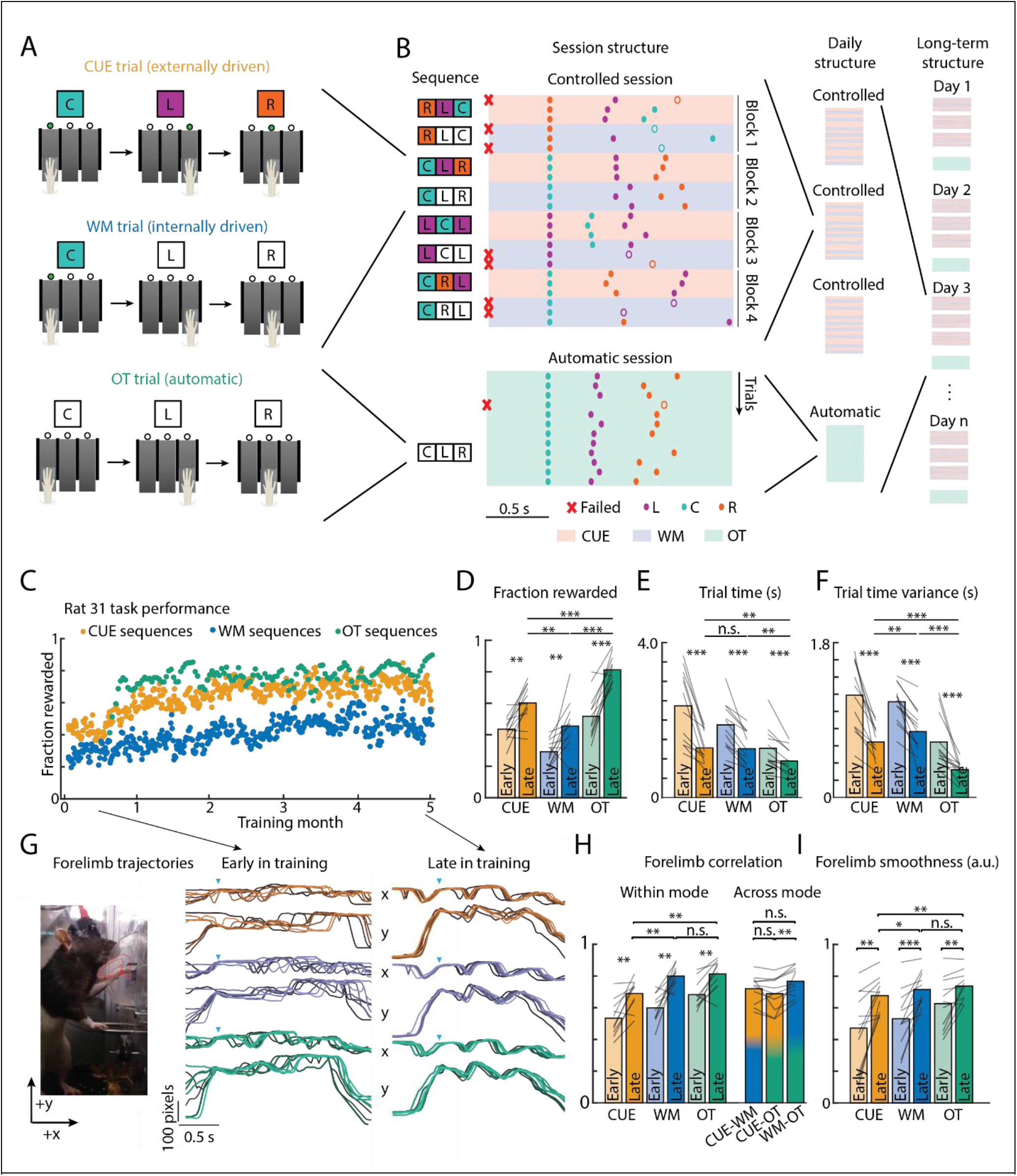
Paradigm for training rats to produce visually guided, working memory guided, and automatic motor sequences. **A**. Rats learn to generate 3-element lever-press sequences that, in ‘controlled’ training sessions, are either visually cued (CUE) or generated from working memory (WM) by repeating the sequence from the preceding cued trial. In a separate ‘automatic’ session without any cues, the same sequence is rewarded each day. The overtrained (OT) sequence is chosen randomly for each rat and fixed for the duration of the experiment. **B**. Performance of an example rat in a controlled (top) and automatic session (bottom). In controlled sessions, sequences are randomly chosen, from 12 possible sequences, and change every block. **C**. Fraction of rewarded trials per session in the three execution modes (orange = CUE, blue = WM, green = OT) for one animal over several months of training. **D-F**. Performance metrics improve from early (light shade) to late (dark shade) in learning across all three execution modes (n=12 rats). **D**. Fraction of rewarded trials. **E**, Trial time (middle). **F**. Variance in trial time (right). **G**. Vertical (y) and horizontal (x) components of forelimb trajectories, aligned to the first lever press, for 8 example trials from early and late in learning, as captured from a side camera. Examples are selected from trials with similar duration. **F**. Trial-to-trial correlation of the active forelimb trajectory within a given execution mode from early and late in training (n=10 of 12 rats were tracked in early learning), and correlation of the kinematics across the three modes, late in training. **I**. The smoothness of the movement kinematics, as measured through the inverse of the spectral arc length^71^ (see Methods) increases over learning. Values closer to 1 indicate smoother movements. *P<0.05, **P<0.01, ***P<0.001, Wilcoxon sign rank test.

Rats were initially trained to press a single lever for a water reward, then to associate a visual cue above each lever with pressing that lever (see Methods). After acquiring the cue-action association for each of the levers (5164 ± 948 trials; mean ± s.e.m), rats transitioned from single presses to two- and, ultimately, three-element sequences. In ‘controlled’ training sessions, the rewarded keypress sequence was either signaled directly and sequentially by visual cues (**CUE**) or had to be remembered from the instructed sequence of previous trials (working memory, **WM**; see Methods). This trial design, adopted from studies in NHPs^30^, allowed us to compare performance and neural dynamics for sequences externally guided by visual cues and internally generated from working memory^20,21,76–78^. Blocks of CUE and WM trials were interleaved, and, in each, one of the 12 possible sequences was randomly selected and rewarded (Fig. 2b).

In separate ‘automatic’ sessions (see Methods), animals were trained to produce the very same pre-determined keypress sequence - randomly chosen for each rat from the 12 possible sequences - for the duration of the months-long experiment (overtrained mode, **OT**). Because this automatic sequence is one of the 12 sequences rewarded in the controlled sessions, we could compare the same motor sequence across the three distinct execution modes.

### Rats master the ‘piano playing’ task

Rats learned to produce the prescribed sequences under all three conditions (CUE, WM, OT) (Fig. 2c-d). We deemed rats to be ‘experts’ when both their success rates and trial times were reliably within 0.5 σ of asymptotic performance values (see Methods). Expert performance was reached after 17,623 ± 7616 trials (75 ± 23 days) in the CUE mode, 11330 ± 5278 trials (88±30) days in the WM mode, and 14061 ± 7082 (72 ± 21 days) trials in the OT mode.

The success rate of expert rats was, on an average, 60.19% ± 10.73% (CUE), 44.29 ± 15.45% (WM) and 79.57 ± 11.84% (OT). Note that in all cases, this is many times chance performance, which would be 8.33% considering only the 12 prescribed sequences or 3.7% considering all possible three element keypress sequences. As is expected from similar learning paradigms in humans and NHPs^6,7,20^, the mean and variability of the trial times decreased with learning (Fig. 2e-f), while the stereotypy of the associated movement patterns increased across all three execution modes (Fig. 2g). Finally, to estimate the efficiency, or smoothness^8,79^, of a movement we used the inverse of the spectral arc length^71^, a noise-robust and dimensionless metric that reflects the smoothness fluidity of a trajectory (see Methods). Consistent with expectations from humans and NHPs^8,80,81^, this metric increased over learning.

### Kinematic similarities across execution modes and movement elements

Probing a distinction in how neural circuits implement the different types of motor sequences requires dissociating differences in execution mode from differences in movement kinematics. To establish the degree to which kinematics for the same motor sequence across execution modes (i.e. CUE, WM or OT) were similar, we tracked the rat’s dominant forelimb (i.e. the one pressing the lever) and nose from videos recorded from the side and the top (Fig. 2g, Supp vid 1, Extended Data Fig. 1a)^82,83^. Comparing trials of similar duration across execution modes revealed very similar forelimb trajectories. Pairwise correlations of trajectories, linearly warped to lever press times, had similar distributions whether the trials were from within or across modes (Fig. 2h). Nose trajectories were also similar across trials and execution modes and had high pairwise correlations (Extended Data Fig. 1b-c). The similarity in kinematics is important because it allows us to interpret any potential differences in neural activity and sensitivity to neural circuit manipulations as being due to differences in execution mode (OT, WM, CUE) rather than in low-level aspects of motor implementation.

### Overtrained motor sequences have signatures of automaticity

In humans, motor automaticity is distinguished by improved performance, increased movement speed, and less variable execution times^1,11,12^. While we found that the kinematics for the same motor sequence across session types was similar (Fig. 2e-f, Supplementary Movie 1), we parsed these metrics in the behavioral data to assess whether automaticity had been established in OT sessions (Fig 3a-d).

**Figure 3:**
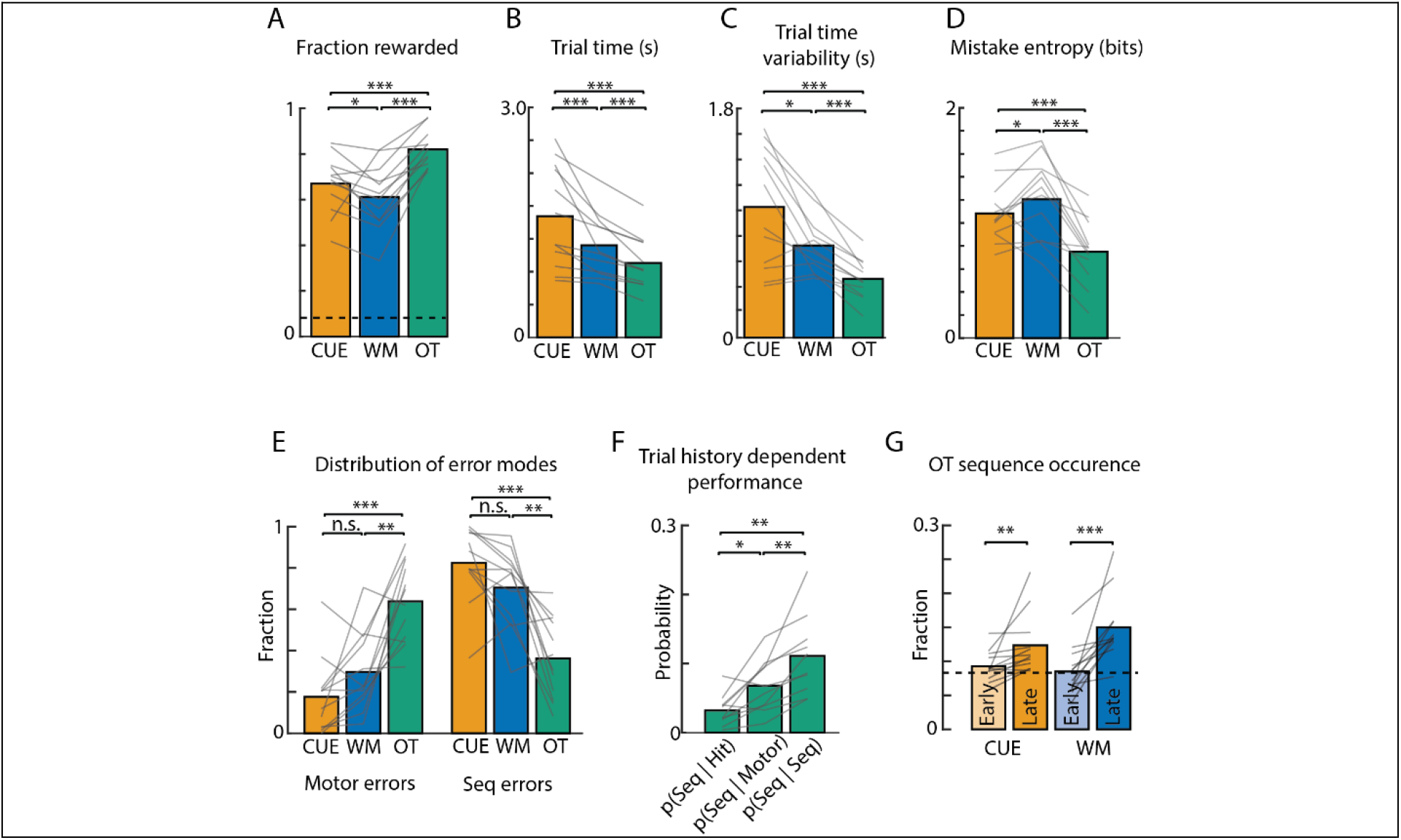
Overtrained motor sequences show signatures of automaticity. **A-D**. Performance metrics for the same sequence across execution modes. Gray lines are averages within rats; bars are the grand average across rats. **A**. Success rate, or fraction of rewarded trials. **B**. Median trial duration for rewarded sequences measured as the time from the 1^st^ to the 3^rd^ lever press. **C**. Standard deviation of trial durations. **D**. Variability, or Shannon entropy, of the mistakes made in unrewarded trials (see Methods). **E**. Fraction of unrewarded trials classified as either motor errors or sequence errors (see Supplementary Movie 2 and Methods). **F**. Probability of a sequence error, if the previous trial was rewarded (Hit), a motor error, or a sequence error (Seq), in the OT condition. **G**. Probability that the rat performs the overtrained sequence in the controlled session before and after the overtrained sessions are introduced. Dotted line is chance of performing any particular sequence (8.33%). *P<0.05, **P<0.01, ***P<0.001, Wilcoxon sign rank test.

Consistent with signatures of automaticity, we found that trials in OT sessions were more successful, faster, and less variable than when performing the same sequence in controlled sessions (Fig. 3a-c). We also examined the failure modes, hypothesizing that rats performing automatic sequences should be more systematic, or less variable, in the errors they make^84^. In agreement with this, we found that the entropy, or randomness, of erroneous sequences was much lower in the OT sessions than for either of the controlled sessions (CUE, WM; Fig. 3d).

Furthermore, if automatic motor sequences are consolidated and defined in terms of continuous low-level motor commands^24,62,63,85^, and not as serial discrete action selection, unsuccessful trials in OT sessions should be dominated by errors relating to variability in movement kinematics (e.g. a forelimb swipe at the ‘correct’ lever that misses the target), as opposed to errors in higher-level sequencing aspects (Supplementary Movie 2).

In support of this, failures in OT trials were mostly due to rats swiping at but missing the ‘correct’ lever or failing to depress it beyond the threshold for detection (approximately 63.81 ± 19.56% of all OT errors were of this type; Fig. 3e). This was in contrast to controlled trials, in which unsuccessful attempts were dominated by true sequence errors, where rats orient towards and press the ‘wrong’ lever (only 17.6 ± 18.25 % of CUE and 29.62 ± 19.02% of WM errors were motor errors). Interestingly, potential sequence errors in the OT sessions were twice as likely to come after trials with motor execution errors compared to correct trials (Fig. 3f, see Methods), consistent with a drop in reward triggering increased motor exploration^84^.

Similar to studies in humans and NHPs performing both sensory cued and automatic motor sequences^4,48,86^, rats were more likely to execute the overtrained sequence in controlled sessions compared to chance (12.35±4.43% in CUE, 14.99±4.85% in WM, where chance is 8.3%; Fig. 3g).

These differences in the quality of motor sequence execution and the error modes across the distinct session types are consistent with studies comparing sensory cued and automatic motor sequences in humans^6,7,13^ and indicate that automaticity of the overtrained motor sequence, as commonly defined in the literature, had been achieved.

### DLS represents motor sequences similarly irrespective of execution mode

We had designed our behavioral paradigm to directly probe whether and how the neural implementations of different types of motor sequences differ at the level of the striatum. If its sensorimotor region, DLS in rodents, distinguishes modes of execution at the action selection-level, as widely believed^43,44,46,47,58^, we argued that it should manifest as a difference in the associated neural activity patterns between visually-guided and automatic execution modes (Fig 1). Neural recordings should also speak to whether and how the BG’s role in high-level action selection differ across visually- and working memory-guided sequences.

Alternatively, if BG encodes sequences at a ‘lower’ execution level (Fig. 1), we may see no difference in the activity patterns, as the motor outputs are kinematically similar across modes (Figs. 2g,h). To directly and effectively probe this, we implanted 64-channel tetrode drives targeting the DLS in expert rats (n=4, Extended Data Fig. 2a) and compared the activity of the same neurons for the same motor sequence in OT, CUE and WM trials. We recorded neural activity continuously over several weeks, comparing units recorded for at least 5 trials for each execution mode (CUE, WM, OT) for the same motor sequence (in total: n=579 neurons selected from 2468 total, see Methods).

The comparisons across each execution mode (Fig. 4a) were striking for the lack of any qualitative difference: task-related activity patterns of DLS neurons for the same motor sequence were highly correlated across all trial types (CUE, WM and OT; Fig. 3a-b,d) with no significant difference in either the average firing rates or peak z-scored activity (Fig. 4c). We found a similar result when splitting the population into putative medium spiny neurons (MSNs) and fast-spiking interneurons (FSIs) (Extended Data Fig. 3). Thus, the neural recordings, on their own, did not suggest a meaningful difference in how the BG are engaged in sensory-guided, working memory-guided, and automatic motor execution despite prior suggestions to the contrary^15,19,37–39,49,51^.

**Figure 4:**
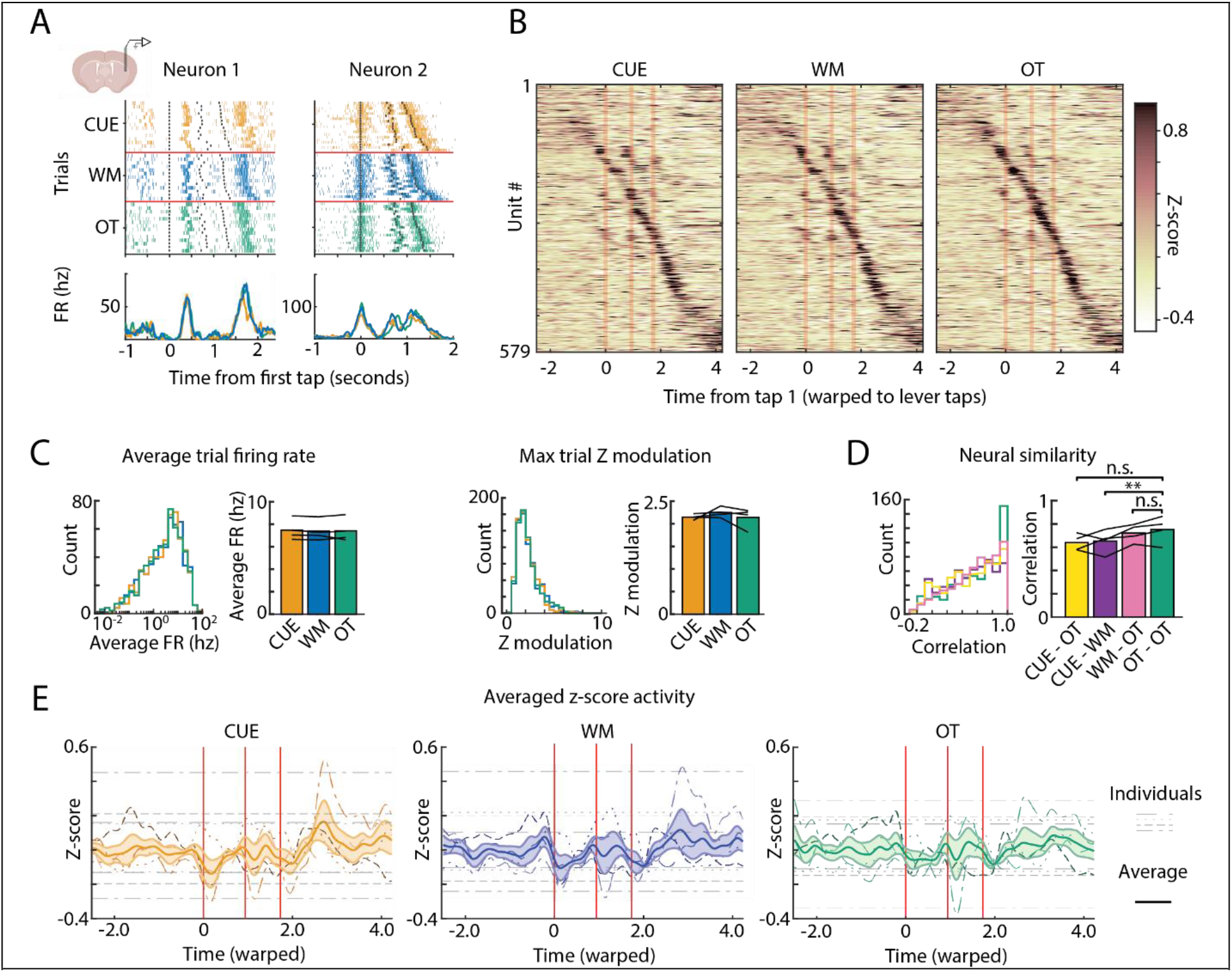
DLS represents motor sequences similarly across execution modes. **A**. Spike rasters for two example neurons recorded in DLS during the execution of the same motor sequence in OT (green), CUE (red), and WM (blue) trials. Black dots indicate the time of a lever-press; red lines separate the execution modes. Below is the instantaneous firing rate across each execution mode. Trials were subsampled to have equal trial durations across execution modes. **B**. Z-scored average activity of 579 neurons recorded in the DLS during execution of successful OT, CUE, and WM trials with the same target sequence (from n=4 rats). The trials were linearly time-warped to each lever press (red vertical lines). Units were sorted by the time of their peak activity. The sorting index was calculated from half the available trials for each unit, taken from the OT mode, and then applied to the remaining trials and modes. **C**. Comparing task-aligned activity statistics across the population of recorded neurons. (Left) Histogram of the average firing rate during the trial-period (P>0.05, paired t-test for each execution mode). The bar graph breaks this down by rats (n=4). (Right) Same as in the left panel but for maximum modulation of Z-scored firing rate during the trial-period (P=.94 OT-CUE, P=.25 OT-WM, P=.26 CUE-WM, paired t-test). **D**. Histogram of correlation coefficients between trial-averaged activity across execution modes for the DLS neurons shown in **B** (see Methods). Left, histogram of the correlation coefficients of the trial-averages neural activity on CUE and OT trials (yellow), CUE and WM trials (purple), WM and OT trials (cyan), and OT and a heldout set of OT trials (green). Right, mean correlation coefficients for trial-averaged activity across execution modes broken down by animal (n=4). **P<0.01, paired t-test. **E**. *Z*-scored firing rates averaged over all neurons. Thick, solid lines indicate the grand average across all rats (n=4), and colored shaded regions indicate the s.e.m. across rats. Thin, dashed lines indicate individual rats, and the thin, dashed gray lines denote the 95% confidence interval of z-scored activity. Average firing rates are highly correlated across all execution modes for each rat (0.8286 +-0.0874, mean +-s.e.m) and do not significantly differ at the time of the first lever press (p>0.05, two-sided t-test, n=4 rats).

### The DLS does not encode high-level features of the sequence

We next analyzed neural activity patterns in DLS for clues about the general contributions it makes to motor sequence execution, focusing first on its putative role in discrete action selection. Prior studies supporting such a function have interpreted elevated average DLS activity at the boundaries of ‘chunked’ actions^52,55,56,87^ as an indication that the BG help bias their initiation and/or termination by facilitating and/or inhibiting downstream control circuits^58^. To test whether activity is concentrated at the boundaries of putative action ‘chunks’, we examined the firing rate modulation over the length of the whole sequence (Fig. 4e; see methods). We found no evidence for population activity in the DLS preferentially marking the start and/or stop of overtrained sequences; neither did it consistently mark the boundaries between behavioral elements (lever-presses, orienting movements) or pairs of such (combined orient and lever-press movements) (Fig. 4e).

While average population activity in DLS did not demarcate the boundaries of motor elements or motor chunks, its activity could reflect their place in the sequence in other ways^46,47,88^. For example, it has been proposed that DLS neurons represent the sequential context of movements and actions, including their ordinal position or, in lever-pressing tasks, the identity of the lever being pressed^54,89^. In such a coding scheme, DLS activity associated with a specific movement should not be a mere function of its kinematics, but also reflect higher-order features of the sequence^32^.

To explicitly probe this, we expanded our analysis to all 12 motor sequences generated in controlled sessions. If DLS represents higher-order features of sequence organization, its neural activity should differ when the same lever-press or orienting movement is performed in different sequential contexts (e.g., the press and orienting movement L→C in the sequence L→C→L vs. the sequence C→L→C; Fig. 5a). We did not find this to be the case: neural activity across two sequences composed of different elements (L→C→L vs. C→R→C) but similar lever-press and orienting movements (press→move right→press→move left→press), was similar (Fig. 5a). More generally, DLS activity associated with a given motor element was highly correlated regardless of the sequence in which it was embedded, its ordinal position in the sequence (1^st^, 2^nd^, 3^rd^ press), or the specific lever being pressed (i.e. L, C, or R) (Fig. 5a-e). There was however a very clear distinction in how striatal neurons represented orienting movements to the left and right, both short (e.g., L→C) or long (e.g., L→R), consistent with an egocentric kinematic code (Fig. 5d-e, Extended Data Fig. 4a-b).

**Figure 5:**
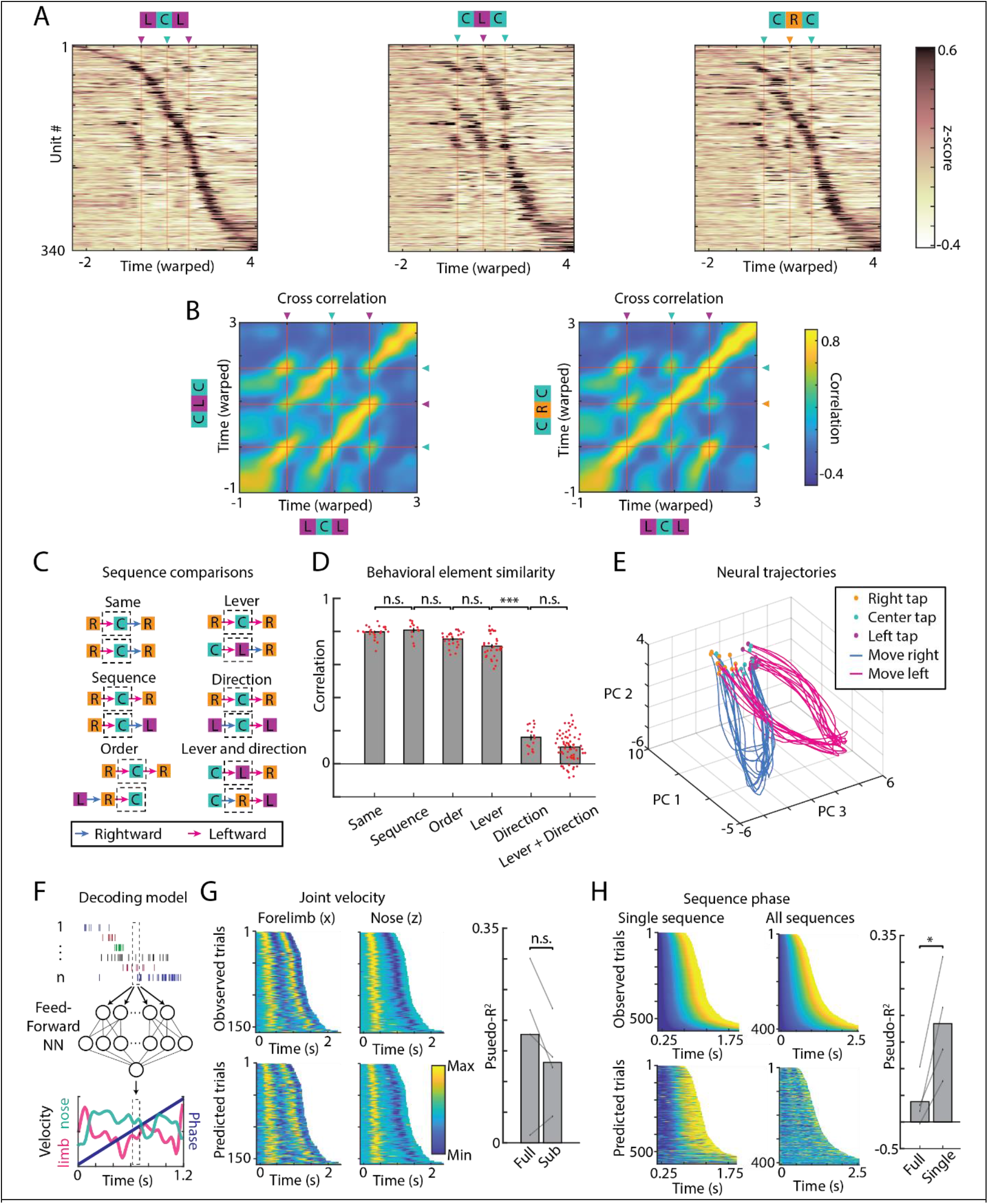
DLS encodes low-level kinematics, not high-level attributes, of discrete motor sequences. **A**. Comparing trial-averaged neural activity for different sequences, from all DLS neurons that met our criteria for inclusion (See Methods), across 4 rats. Units were sorted based on the time of peak firing rate in the LCL sequence (left most). Sequences LCL and CLC (left and center), are comprised of the same motor elements (LC and CL occur in both) but ordered differently. Sequences LCL (left) and CRC (right) are comprised of the same actions (press, move right, press, move left, press), but on different levers. **B**. Cross-correlations of time-varying neural activity from A (see Methods). **C**. The controlled condition allows us to compare the same sub-sequence of orienting and pressing movement (here: rightward orienting and pressing) in different sequential contexts. Dashed box indicates examples of comparisons made in D. **D**. Correlation of the neural activity patterns associated with the same sub-sequence of orienting and pressing movements in different sequential contexts. ***P<0.001, Bonferroni corrected Wilcoxon rank sum test. **E**. Neural population trajectories in PC space for the 12 different sequences, plotted along the 1^st^, 2^nd^, and 4^th^ PC. In DLS, all lever-press movements (color dots) are associated with similar neural activities, as are all right-(cyan) and left-ward (magenta) movements, independent of lever identity or order in the sequence (i.e., right→center, center→left, and right→left are all similar). **F-H**. Decoding analysis. **F**: Schematic of the decoding analysis. A feedforward neural network was trained to predict either the instantaneous velocity of the active forelimb (viewed from the side) and the nose (viewed from above), or the sequence phase, from the spiking activity of groups of simultaneously recorded DLS units. **G**: Velocity of the active forelimb (top left) and nose (top right) from a representative session, in the +x and +z dimension respectively (see Methods). Trials are aligned to the first lever press and sorted by trial duration. Velocity predictions on held-out trials for forelimb (bottom left) and nose (bottom right). (Right) Model performance (measured in pseudo-R^2^, see Methods), tested on held-out trials, when predicting all velocity components (forelimb +x, +y, and nose +x, +z, see Methods). Performance is shown for models trained on every sequence (Full) and tested on held-out trials, and for models trained on a randomly selected half of the sequences and tested on the other half (Subset). Gray lines indicate model performance within individual rats (n=4). **H**. Heatmaps show observed (top) and predicted (bottom) sequence phase for single sequences without repeated elements (left), and for all controlled sequences (right). Trials are aligned to the first tap. (Right) Model performance, tested on held-out trials and quantified by the pseudo-R^2^, for models trained on all controlled sequences (full), and those trained on a single sequence with non-repeating elements (single). *P<0.05, two-sided t-test.

### DLS encodes detailed movement kinematics

Based on this initial analysis and related studies^24,60,62,63^, a plausible alternative to DLS representing higher-order aspects of discrete motor sequences is that is encodes - and contributes to shaping - the detailed kinematics of learned sequential movement patterns, including their vigor. While DLS is not required for species-typical lever-press or orienting movements^24,36,75,90^, it can, by acting on downstream control circuits, help make them more adapted to a specific task^24,62,91^ (Fig. 2e). In this scenario, we would expect DLS neurons to encode kinematic features continuously throughout the behavior^24,63,85^.

To probe this idea^24^, we trained a multilayer neural network to predict, using the spiking activity of simultaneously recorded DLS units, the instantaneous velocity of the rats’ active forelimb and nose during the controlled task, as viewed from a side and top camera respectively (Fig. 5f, Extended Data Fig 5a). Consistent with observations from trial-averaged ensemble activity, we could decode movement kinematics on individual trials across the different sequences from populations of DLS neurons (Fig. 5g). Decoders trained on a subset of sequences could predict kinematics from held-out sequences just as well (Fig. 5g), implying that the kinematic code in DLS is invariant to the sequential context of the movements as suggested by our earlier analysis (Fig. 5d-e). Finally, to determine if we could decode the 3D kinematics beyond what is captured in the 2D camera projections, we triangulated the forelimb and nose position into 3D using multiple views across our three cameras (Extended Data Fig 5a-c, Methods). Decoders of DLS activity could predict variations in movement kinematics on individual trials similarly in 3 dimensions (Extended Data Fig 5d-f).

Recent studies^85,92,93^, have suggested that DLS encodes the progression, or the ‘phase’, of a behavioral sequence. In the context of stereotyped movement sequences, however, kinematics and phase are tightly coupled, making it difficult to parse which of these attributes DLS activity reflects^94–96^. Because our controlled task condition breaks this coupling, kinematics and phase become dissociable. Training a decoder to predict the phase of the behavior, however, failed (Fig. 5h), meaning that phase in the sequence cannot be recovered from DLS activity alone. Only when phase and kinematics was coupled, i.e. when we considered only single sequences with no repeated motor elements (e.g., lever taps, see Methods for how these were selected), could we decode phase (Fig. 5h). Taken together, our results suggest that DLS encodes low-level continuous kinematics of movements in a way that is - in contrast to prior reports^54,56^ – invariant to their sequential context.

### Probing DLS function by lesions

Although our neural recordings showed that DLS activity reflects ongoing kinematics in a similar way for sensory-guided, working memory-guided, and automatic motor sequences, this does not establish a causal role for DLS in their execution. Alternatively, DLS activity could – in one or all modes – simply reflect input from essential sensorimotor control circuits^97^. To arbitrate between these possibilities, we lesioned DLS bilaterally in expert animals (n=7 rats, see Methods, Fig. 6a, Extended Data Fig. 2b).

**Figure 6:**
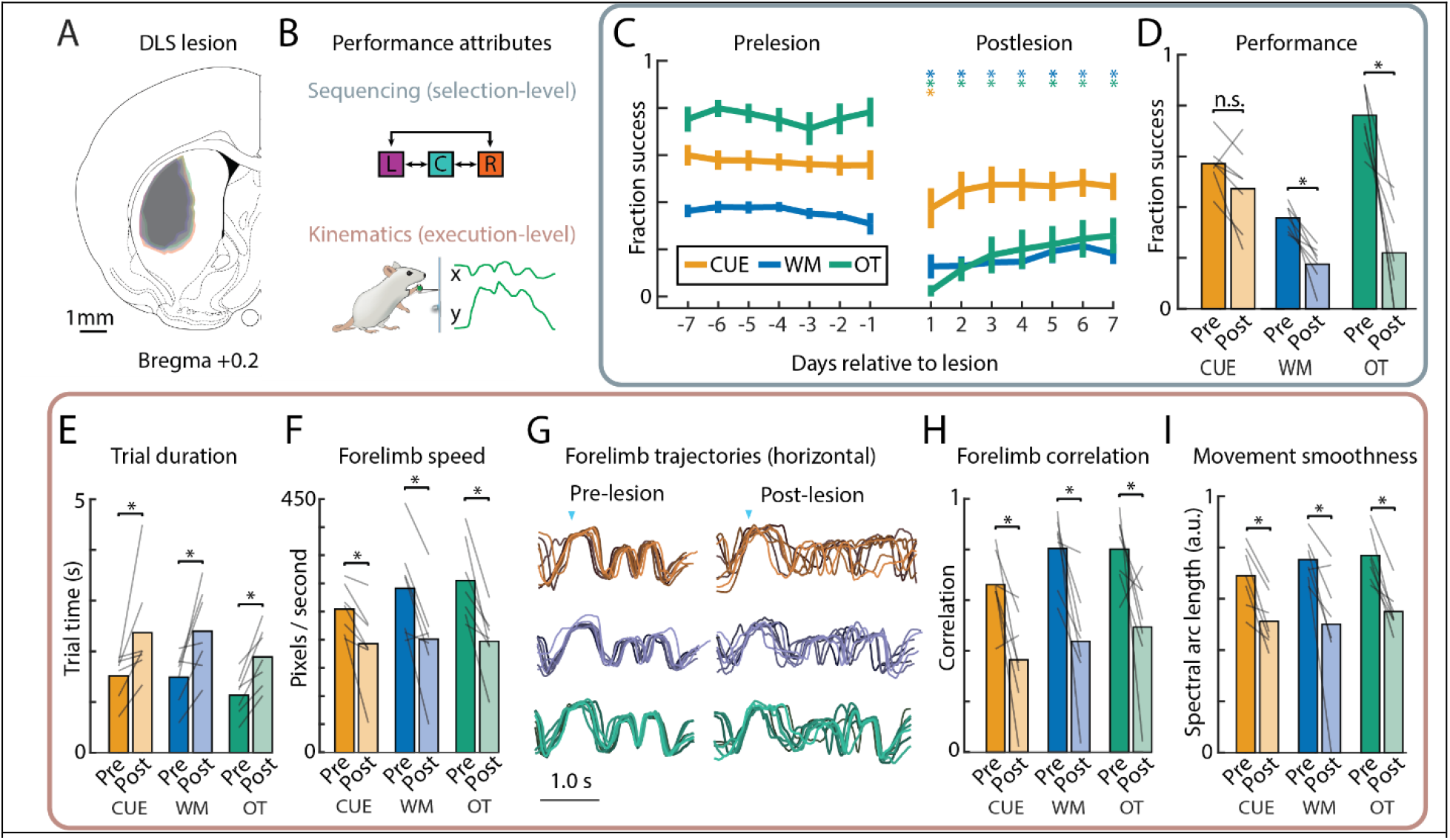
DLS is required for generating task-specific movement kinematics across all modes, but is not required for ordering basic movements in the prescribed sequence when visually-cued. **A**. Outline of DLS lesion boundaries, from 3 rats in one hemisphere. For full lesion annotation, see Extended Data Fig. 2b. **B**. Schematic of the two performance attributes we parse. The first is high-level sequencing of orienting and pressing movements. Success (reward) in the task is contingent on getting this right. The second is the low-level implementation of the requisite movements, i.e task-specific learned kinematics. While movements adapted to the task are faster and more fluid than at the onset of training, they are not required for reward. **C-D**. Effects of DLS lesions on sequencing. **C**. Success rate in producing the prescribed sequence, averaged over rats (n=7, error bars are SEM), in the week before and after lesion. Stars denote whether performance is significantly different on a given day, relative to average performance in the week pre-lesion, for each execution mode. **D**. Bars show success rate in the week pre-lesion and days 3-7 post-lesion, averaged across rats. Gray lines denote individual rats. **E-I**. Effects of DLS lesions on movement kinematics. **E**. Trial times and **F**. average forelimb speed for trials before and after the lesions (see Methods). **G**. The horizontal (x) position of the active forelimb on eight example trials with similar duration are overlayed and compared before (left) and after (right) DLS lesions. **H**. Average trial-to-trial correlation and **I**. movement smoothness for trajectories of the active forelimb (both horizontal (x) and vertical (y)) before and after lesion. *P<0.05, **P<0.01, Wilcoxon sign-rank test.

In interpreting the effects of striatal lesions, we distinguish, as we did for the neural analysis, two aspects of performance, each associated with a putative function of the BG (Fig. 6b). The first is the ability to perform the prescribed sequence of lever presses (i.e. ‘sequencing’). The second is the ability to use fast and efficient movements refined and adapted to the task (i.e. ‘kinematics’). Note that only the sequencing aspect is required for reward. Parsing performance in this way allows us to probe whether striatum contributes to high-level sequence structure and low-level movement kinematics differently across execution modes.

### DLS lesions affect high-level sequence structure on automatic and working-memory, but not visually cued, trials

Following a seven-day post-lesion recovery (see Methods), the rats’ ability to perform the OT sequence was severely impaired (Fig. 6c-d), dropping to near chance levels (8.33%). In stark contrast, success rates on CUE sequences were comparable to pre-lesion (Fig. 6c-d), save for a brief drop on the first few sessions after the lesions consistent with non-specific transient of the surgery^24,25^. Success rates in WM trials, on the other hand, were chronically affected, and while the relative drop was less than for the OT mode, performance still dropped to near chance levels (Fig. 6c-d). A seven-day control break before the lesion (see Methods), did not significantly affect the behavior in either execution mode (Extended Data Fig. 6).

One possible explanation for the lesion resilience in CUE trials, and the less severe performance drop in WM trials (compared to OT), is that visual cues aid movement initiation in a DLS-independent manner^90^. However, the post-lesion drop in success rate on OT and WM trials could not be explained by a deficit in movement initiation^62^. Although DLS lesioned rats generated fewer lever-presses overall, they were actively engaged in the task and performed similar numbers of trials across controlled and automatic sessions (Extended Data Fig. 8a).

Taken together, these results are consistent with DLS having an essential role in controlling high-level sequence structure for automatic (OT) and working memory-guided motor sequences (WM)^24,49,98^. However, we found DLS to be dispensable for sequencing visually cued behaviors (CUE)^99,100^.

### DLS lesions affect learned movement kinematics equally across execution modes

We next probed the effects of DLS lesions on movement kinematics, i.e. the aspect of the animals’ performance we could reliably decode from DLS activity. Consistent with a general role for the BG in the specifying learned task-specific movement kinematics^24^, orienting and lever-pressing movements across all three execution modes were similarly affected. The ‘vigor’ of the movements, defined as the scalar gain factor applied to the kinematic features of a movement such as movement latency or speed^101,102^, was reduced (i.e. longer trial times and slower speeds, Fig. 6e-f). The change in vigor following lesion was similar for short (e.g. L->C) and long (e.g. L->R) movements (Extended Data Fig. 7a-d).

One plausible coupling between deficits in kinematics/vigor and the ability to generate the proper sequence is if the neural dynamics that inform the sequence in OT and WM trials can only be expressed at higher (i.e. closer to pre-lesion) speeds (CUE trials would be paced and informed by visual cues and hence would not be affected). However, that we did not find a consistent relationship between vigor (movement speed) and trial outcome after DLS lesions (Extended Data Fig. 7e-h), speaks against this idea.

While the lesion-induced effects on overall movement speed and latency are consistent with prior reports on BG’s role in controlling movement vigor^48,60^, other aspects of kinematics were also affected. Trial-to-trial movement variability, even for similar duration movements, was dramatically increased (Fig. 6g-h). Furthermore, the smoothness of task-related movements, which increases over learning^71^, also decreased following DLS lesions, a finding that mirrors what is seen in Parkinson’s patients^103^ (Fig. 6i). Interestingly, we found that the quality of the movements post-lesion reverted to what is seen early in learning (Extended Data Fig. 8b-g), a result consistent with DLS learning to control task-specific movement kinematics^24,26^ by interacting with downstream control circuits that generate species-typical movements^23^.

### The DMS is not required for motor sequence execution in either mode

Our focus on sensorimotor striatum (DLS) was motivated by its known role in movement execution^43,61,91^. In contrast to DLS, which receives much of its input from sensorimotor cortex, dorsomedial striatum (DMS) receives its cortical input predominantly from PFC and PPC^104,105^. Though this associative region of the striatum has been implicated in flexible control of behavior, such as modifying, switching, and updating behavioral choices in response to previously learned associations^65–68,106^, whether it has an essential role in generating sensory- or working memory-guided motor sequences is less clear^36,75^. To probe this, we lesioned DMS bilaterally in a separate cohort of expert animals (n=6 rats, see Methods, Fig. 7a, Extended Data Fig. 2c).

**Figure 7:**
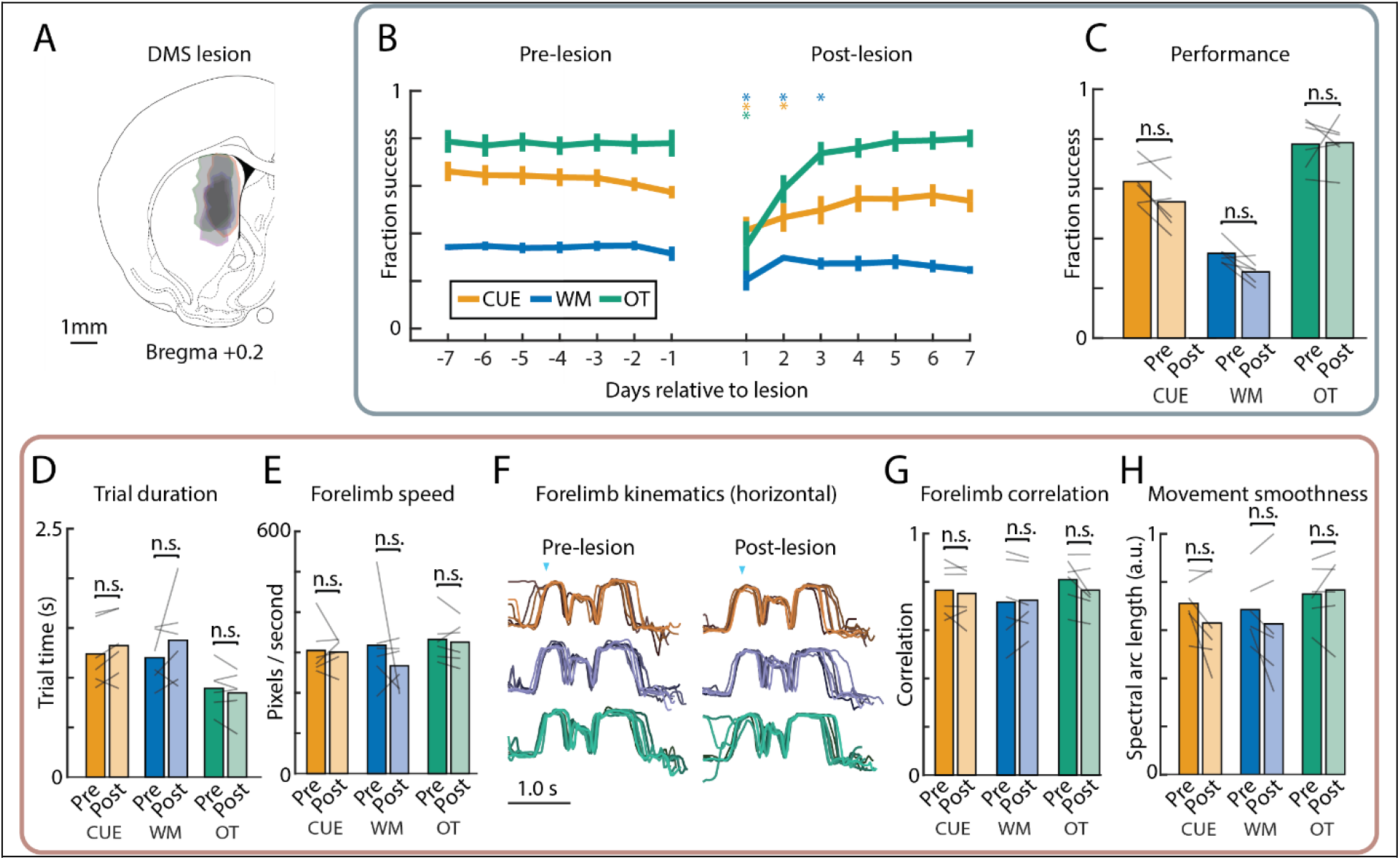
DMS lesions have no long-term effect on either controlled or automatic sequence execution. **A**. Outline of DMS lesion boundaries, from 3 rats in one hemisphere. For full lesion annotation, see Extended Data Fig 2c. **B-C**. Effects of DMS lesions on sequencing. **B**. Success rates in producing the prescribed sequence, averaged over rats (n=6, error bars are SEM), in the week before and after lesion. Stars denote whether performance is significantly different on a given day, relative to average performance in the week pre-lesion, for each execution mode. **C**. Bars show success rate in the week pre-lesion and days 3-7 post-lesion, averaged across rats. Gray lines denote individual rats. **D-H**. Effects of DMS lesions on movement kinematics. **D**. Trial times and **E**. average forelimb speed before and after the lesions (see Methods). **F**. The horizontal (x) position of the active forelimb on eight example trials with similar durations are overlayed and compared shown before (left) and after (right) DMS lesions. **G**. Average trial-to-trial correlation and **H**. movement smoothness for trajectories of the active forelimb (both horizontal (x) and vertical (y)), before and after lesion. P>0.05, Wilcoxon sign-rank test.

Consistent with our previous work^24^, we found that DMS was not required for automatic motor sequence execution (Fig. 7b-c). More surprisingly, DMS lesions also did not have any lasting effects on controlled motor sequence execution in either WM or CUE trials (Fig. 7d-h). While there was a transient decrease in success rate in the first few days following the lesion, this is consistent with nonspecific effects of the surgery procedure and recovery (see similar effects in^24–26^ and Fig. 6c - CUE condition). While we cannot rule out that these transient effects reflect a real contribution of DMS to motor sequence execution, we note that any such putative function can be compensated for in a matter of days. Furthermore, unlike for DLS, lesions of DMS did not significantly affect kinematics in either execution mode, with lesioned rats showing no consistent increase in trial time or mean trial speed or a drop in the stereotypy of their task-related movements (Fig. 7d-h). This reinforces the dissociation between the DLS and DMS in terms of low-level kinematic control^24^, and further suggests that DMS is not necessary for either sequencing or kinematic aspects of well-trained motor sequences. Whether DMS plays a role in early motor sequence learning remains an intriguing open question^34–36,87,107^.

### A simple neural network model can account for the results in both CUE and OT execution modes

At first glance, our results suggesting both similar (e.g., in terms of coding properties and contribution to low-level kinematics) and different (e.g., in terms of effects of lesions on high-level sequence structure) functions for DLS across modes, may seem discrepant. To reconcile these findings and better inform the circuit-level logic underlying motor sequence execution, we built a simple neural network model of the motor system (Fig. 8a) in which a DLS-like circuit learns to interact with ‘downstream’ control circuits in different execution modes. Our modeling focused on sensory-guided and automatic execution modes, as these show the clearest distinctions in our experimental data, setting aside for this analysis the working memory-driven condition.

**Figure 8:**
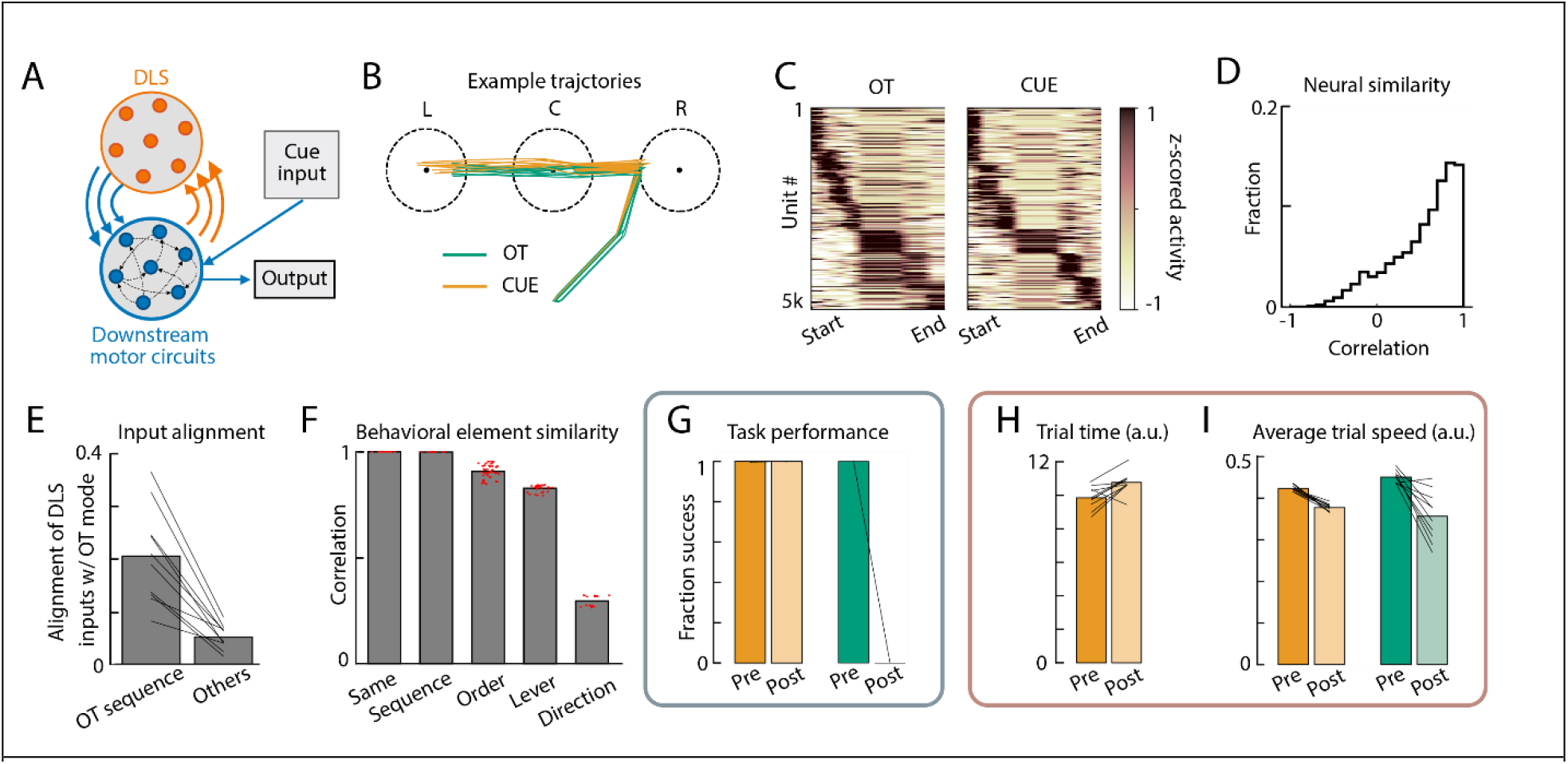
Experimental results emerge naturally during task learning in a dual module neural network model. **A**. Schematic illustrating the architecture of our neural network model. DLS weights (orange) are trained on the cued and automatic task modes and interact with downstream recurrent motor circuits already trained to perform cued movements. **B**. Example trajectories on the simulated task. The network controls the velocity of a ‘forelimb’ and must move it into three circular regions (representing ‘level-presses’) in the correct sequential order (in this example, RCL for both the CUE and OT modes). **C**. Average Z-scored activity of model neurons in the DLS population during execution of successful OT and cued trials with the same target sequence. Neurons are sorted by the time of their peak activity, and displayed in the same order in both plots. **D**. Histogram of correlation coefficients between CUE and OT modes over all model DLS neurons of correlation, as in Fig. 4d. **E**. Correlation between input to DLS units across modes. Line segments indicate individual simulations; bars indicate averages cross simulations. Left: correlation between DLS inputs on OT and cued trials for the same sequence. Right: same measurement but shown for cued trials with other target sequences (averaged over all other sequences). **F**. Correlation of the neural activity patterns associated with the same sub-sequence of orienting and pressing movements in different sequential contexts, as in Fig. 5d. **G**. Effects of DLS removal on sequencing. Average success rate in producing the prescribed sequence., averaged across ten network simulations for each condition. Lines depict individual simulations. **H-I**. Effects of DLS removal on kinematics. **H**. Trial time (i.e., number of simulation steps) and **J**. trial speed (average magnitude of velocity, in arbitrary simulation units) averaged across ten network simulations for each condition.

The DLS network in our model contained no recurrence, reflecting the fact that spiny projection neurons are coupled via relatively weak and sparse lateral inhibition^108,109^. The ‘downstream’ component of the model – intended to capture the control circuits modulated by the BG – was made recurrent. For simplicity, and to focus on the DLS, we abstracted away the details of how it connects to downstream control circuits (i.e., through other BG nuclei, thalamus, etc.), modeling them with a set of linear synaptic weights. To capture the ability of animals to execute cued lever presses prior to sequence training, we pretrained our model to perform a simple in-silico version of ‘lever-pressing’: moving a virtual manipulandum to a target in a 2D environment (Methods).

We then trained the DLS input synapses such that the network’s output ‘moved’ through sequences of three targets (Fig. 8b, Methods). The same network was tasked with producing sensory-guided sequences instructed by external inputs (CUE mode), and also a single internally generated sequence (OT mode; see Methods).

Since animals value time^110^ and reduce trial duration as a function of learning (Fig. 2d), our training procedure incentivized the model to reach the three target positions in the correct order as quickly as possible. The decision to train the DLS input synapses was inspired by the presumed role of cortico-striatal and thalamo-striatal plasticity in reinforcement learning^26,67^.

To compare the performance of our network model to that of real rats, we conducted analyses of the model analogous to those performed on our experimental data. We found that the activity of the neural units in the trained DLS network was largely independent of execution mode as seen in our data (Figs. 8c-d; compare with Figs. 4b,d). This cross-mode independence reflects an emergent alignment of DLS inputs (which are trained on both task modes) for OT and CUE trials with the same target sequence (Fig. 8e). We also recapitulated our experimental observation that DLS network representations reflected egocentric direction-of-motion information (Fig. 8f; compare with Figs. 5d-e).

Next, we simulated lesions to the DLS by removing this part of the model following training, comparing the effects on sequencing and kinematics separately. We found that DLS removal left high-level sequencing impaired for OT but not CUE trials (Fig. 8g; compare with Fig. 6a), recapitulating our experimental results. The model also captured the effect of DLS lesions on movement trajectories, decreasing movement velocity and increasing trial time across execution modes. This is consistent with a role for DLS in adapting movement kinematics for a specific task independent of execution mode (Figs. 8h-i; compare with Figs. 6a-b).

Thus, key features of the experimental data – invariance of DLS activity to execution mode, sensitivity of automatic task performance to DLS lesion, and a role for DLS activity in shaping learned movement kinematics – all emerge naturally during task learning when using a network model with basal ganglia-like circuitry.

In the model, we used a biologically-inspired circuit architecture (Fig. 8a) and matched the training procedure of the network to that of our rats (i.e., with sequence training overlaid on pretrained circuits). To probe how different features of our model contributed to the results, we also considered three alternative circuit models/paradigms. First, to test whether our results depended on DLS output being time-varying and high-dimensional, so as to specify detailed kinematics, we made the DLS output scalar, thus constraining it to represent coarse sequence-level information, e.g. movement speed modulation or ‘vigor’, as has been suggested^48,60^. This model led to markedly different DLS activity patterns across execution modes, in violation of our experimental findings (Extended Data Fig. 9a-g).

We next explored an ‘action selection’ model in which DLS is constrained to generate signals only at the boundaries of elementary movements, which inform the rest of the circuit of the next ‘lever-press’ to be executed. This model resulted in DLS activity patterns that lacked prominent representations of egocentric movement information, again in violation of our experimental data (Extended Data Fig. 9h-m). Finally, to test whether the mode-specific deficits of our lesions were due to the existence of pre-trained motor circuits capable of executing cued movements, we eliminated the pre-training of downstream circuits and trained the full network de novo on all aspects of both task modes (CUE and OT). This simulation failed to capture the DLS lesion resilience seen in our experiments (Extended Data Fig. 9n-s).

Comparing the results from our various models with our experimental data further supports the idea that, for behaviors with learned task-specific kinematics, the BG provide fine-scale kinematic control signals to downstream circuits, which in the case of automatic motor sequences defines the high-level sequential structure of the learned behavior.

## Discussion

We set out to probe the BG’s contribution to motor sequence execution and how it differs depending on whether the sequence is informed by sensory cues, working memory, or is generated after lengthy overtraining (Fig. 1). Surprisingly, neural activity patterns in the sensorimotor striatum (DLS) of rats producing the same three-element sequences were similar across the execution modes (Figs. 2-4). In all cases, DLS activity represented low-level kinematic features – not high-level aspects (e.g. lever identity, ordinal position) – of the behaviors (Fig. 5). Consistent with this coding scheme, lesions to the DLS affected task-specific movement kinematics similarly in all modes. Interestingly, higher-level sequential organization, while not represented in DLS, was affected by DLS lesions on overtrained and working memory-guided trials but remained intact for visually cued sequences (Fig. 6). Lesions to the associative regions of the striatum (DMS) had only transient effects on performance and no effect on movement kinematics (Fig. 7). A simple network model could recapitulate our results and provide an explanation for the different findings: DLS learns to transform its inputs into mode-invariant activity patterns that provide similar kinematic control signals to downstream motor circuits in all execution modes (Fig. 8).

### DLS’s role in specifying low-level kinematics of task-specific learned movements

The BG have been implicated in a diverse array of functions related to motor sequence execution, including action selection^33,44,57,88,111^, the storage of sensorimotor associations^24,97^, chunking^57^, and low-level kinematic specification of the requisite movements^24,48,62,63^. Evidence for the different functions comes from studies that challenge animals to produce motor sequences in different ways^14,112^, some relying on sensory cues others on working memory to inform sequencing^47,49,51,72^, while many probe overtrained, or automatic, sequence execution^24,35,53,56,85^. Thus, the often-discrepant views of BG’s role in motor sequence execution could simply reflect the different computational demands of the various tasks and BG’s ability to contribute to each of them in different ways and to different degrees.

Our experimental paradigm allowed us to test this by probing striatal function across three distinct modes of motor sequence execution in the same animal. Surprisingly, we did not find any meaningful difference in DLS activity when animals performed the same motor sequence in the different execution modes. In all cases, DLS represented ego-centric movement kinematics, suggesting a general role for the BG in the specification and control of learned movements^24,62,63,85,91^.

Most prior studies advancing this view^24,63,85^ probed highly stereotyped movement patterns, in which kinematics and behavioral phase (or time) are intrinsically linked^95^, leaving open the possibility that BG controls not kinematics but the temporal progression^92,96,113^. Our ‘controlled’ motor sequence task breaks this link, allowing us to distinguish the two coding schemes by comparing across many different sequences. This analysis revealed that DLS indeed represents kinematics, not temporal progression, or phase, of the behavior (Fig. 5h).

Consistent with a role for the DLS in specifying time-varying kinematics, lesions affected task-specific learned movements across execution modes (Fig. 6e-f). While lesioned animals could still orient towards the levers and press them, the speed and variability of their movements reverted to what they expressed early in learning (Extended Data Fig. 8b-f). We observed a similar reversion also in a previous study, in which rats learned idiosyncratic and highly precise and complex movement patterns required to press a lever with an inter-press interval of 700ms. After DLS lesions, rats reverted back to species-typical repetitive lever-pressing behavior akin to what they had expressed early in learning^24^.

Together, these results are consistent with a role for DLS in the task-specific refinement of species-typical movements generated in downstream circuits, in our case, likely in the brainstem^114–116^. We saw a similar result in our circuit model: when the ‘DLS’ circuit was allowed to interact with pre-trained ‘downstream circuits’ that could independently respond to cues, variability in trial duration decreased following sequence learning and increased following DLS lesion for both automatic and cued execution modes (Fig. 8h-i). Dissecting the model revealed that, despite receiving different inputs across modes, the DLS learned mode-invariant activity that modulated movement kinematics through its downstream targets.

Our results also corroborated many studies that found DLS lesions to affect movement vigor (the overall speed/amplitude of a movement/action) (Fig 6b-c), a scalar aspect of kinematics^48,60,85,110^. However, vigor alone was not sufficient to explain the change in movement variability and smoothness following DLS lesions (Fig 6e-f), consistent with a role for the DLS in the detailed time-varying specification of movement kinematics^24^. Finally, a model constrained such that DLS provides a vigor signal to downstream circuits showed markedly different activity across execution modes in contrast to our data (Extended Data Fig. 9).

### DLS role in generating high-level sequential structure?

Despite our experiments implicating the BG in the specification of task-specific movement kinematics^24,62,85^, successful performance in our task does not explicitly require such adapted kinematics. Reward is delivered contingent only on the three-element sequence being generated as prescribed, which can be done also with slow and inefficient movements akin to what DLS lesioned animals express. That we nevertheless see deficits in task performance after DLS lesions would seem to implicate the DLS also in the control also of high-level sequence structure.

However, we found no evidence for DLS activity representing such sequence structure. Furthermore, while DLS lesions severely affected performance in automatic and working memory-guided sessions, there was no lasting effect on the success rate on visually cued trials despite the kinematics being similarly affected. Interestingly, this mirrors what is seen in Parkinson’s patients, whose inability to execute internally generated motor sequences can be rescued by providing instructive visual cues^27,117^. Taken together with our results, this implies that cue-action associations, and the serial action-selection process they inform, is not reliant on the BG.

That DLS lesions affected sequence structure on trials in which the prescribed sequence was not informed by external cues (OT, WM trials), raises the question of how the DLS, and the BG more generally, contributes to sequencing such internally generated behaviors? For highly stereotyped overtrained motor sequences (expressed on OT trials), we posit that the behavior becomes consolidated in terms of DLS-dependent continuous low-level kinematics^12,32^. In this case, behavioral progression would no longer rely on the selection of discrete actions but instead results from an invariant and continuous mapping of past to future behavior, a process we argue is DLS dependent^14,24^.

DLS’s contributions to working memory-dependent motor sequences is more of a mystery, though our results suggest that DLS-dependent dynamics is required (Fig 6). One possibility is that striatum is essential for working memory processes more generally. Studies showing that de novo Parkinson’s patients have performance deficits in a working memory task independent of the severity of their motor symptoms supports this idea^98,118^. Though we did not see a distinction in task-related DLS activity between working memory trials and other types of trials (Fig. 4), our study was not designed to assess the time leading up to trial initiation, leaving open the possibility that there may be signatures of working memory processes in this preparatory phase, as has been observed in DMS in a different task paradigm^119^. Interestingly, lesions of DMS, a region implicated in working memory in other studies^120^, did not cause performance deficits in our paradigm (Fig. 7).

### DLS role in ‘chunking’ and action selection?

The BG have also been widely implicated in the process of ‘chunking’^57^, through which a motor sequence initially generated by a serial decision-making process becomes linked, over training, in a way that allows the sequence to be selected and executed as a single action^40,57^. Support for BG’s role in chunking comes from studies showing that the neural representation of task-specific motor sequences in striatum becomes sparser with extended practice^52,56,121^, preferentially marking the boundaries (i.e. start and stop) of overtrained, or ’chunked’, behaviors^52,54,56^. Such sparsification is consistent with a role for the BG in selecting actions elaborated in downstream circuits^52,121,122^. For sensory- and working memory-guided behaviors, action selection would occur at the level of individual elements and hence be more granular, while selection for overtrained behaviors would happen at the level of the whole sequence or ‘chunk’.

Our paradigm allowed us to directly probe whether DLS activity reflects such a function by comparing it for the same motor sequence with similar kinematics in an automatic (selection of the sequence as a chunk) and a controlled (serial selection of discrete motor elements) context (Fig. 1). Not only did we fail to see prominent start/stop activity on automatic trials, but we found that controlled and automatic motor sequences share highly correlated representations in the DLS (Fig. 4).

How should the differences between our results and those showing signatures of chunking and – implicitly – the selection of overtrained sequences by the BG be interpreted? First, most studies implicating BG in chunking tend to compare overall activity early in learning to what is seen in the expert (see, e.g.^19,34,55,87,121^). Any observed activity difference could thus reflect either the process of chunking or changes in movement kinematics, including vigor, that occur as a function of training. For example, one recent study found behaviorally-locked sequential activity patterns in DLS already early in learning, while position- and speed-related activity became more prominent after extensive practice^85^. That we see no difference across automatic and controlled motor sequences suggests that start/stop and sequence-specific activity seen in previous studies is not the consequence of motor chunking per se but may instead reflect learning-related changes in motor output or a shift in how the DLS contributes to it.

Alternatively, the lack of sequence-specific activity in our study could mean that the motor sequence we overtrained failed to coalesce into a single ‘chunk’ due to some peculiarity of our experimental approach. We do not find this plausible, given the clear signatures of automaticity we see (Fig. 3) and the lengthy training times. Furthermore, in an earlier study, in which we trained rats to generate highly stereotyped and stable movement patterns without any need for serial decision making or action selection, we similarly did not see start/stop activity^24^. However, we recognize that absence of evidence is not evidence of absence, and we cannot exclude the existence of sequence-specific activity in cells that we were not recording from.

While our results do not support a role for the BG in initiating consolidated motor chunks elaborated downstream, they are consistent with overtrained motor sequences becoming defined as BG-dependent motor chunks. Thus, rather than selecting them, BG’s role in motor chunking could be in transforming discrete motor sequences into single continuous actions in which the high-level sequential structure of the overtrained behavior is specified by BG-dependent low-level kinematics.

### Circuits controlling sensory-guided motor sequences need to be elucidated

Our finding that visually guided motor sequences can be performed as prescribed after DLS and DMS lesions, begs the question of which circuits control the progression of sensory guided behaviors. Work in humans and NHPs have suggested a role for cortex^17,77,123,124^. Though we have shown that motor cortex is not required for executing highly overtrained automatic behaviors in rodents^25^, it remains an open question whether and how motor cortex contributes to sensory-guided motor sequences and the degree to which its function differs across execution modes. Work on skilled reaching movements, i.e. behaviors crucially instructed by visual input, has implicated motor cortex in their execution^125^. This suggests that the need to respond to environmental cues may render motor cortex a necessary controller. Future experiments will be needed to address the degree to which motor cortex’s contributions to movement control depend on the specific challenges posed by a task, and the condition under which its function require the BG.

## Methods

### Animals

The care and experimental manipulation of all animals were reviewed and approved by the Harvard Institutional Animal Care and Use Committee. Experimental subjects were female Long Evans rats 3- to 8-months at the start of training (Charles River).

### Behavioral apparatus

Animals (n=23 total) were trained on our discrete sequence production task (the ‘piano’ task) in a fully automated home-cage training system^126^. Hardware was controlled by Teensy 3.6 and experiments and videos were recorded by Raspberry Pi 3. Home cage training was done in custom-made behavioral boxes. Boxes were outfitted with three levers spaced approximately 2.5 cm apart, and 14 cm above the floor. Plastic barriers 0.25” thick, 2.3” tall, and of 1” extent were placed between each lever to restrict the postures with which a rat can use their forelimb to press levers. A reward spout for water delivery was placed beneath the center lever. Lever presses were registered when lever displacement reached a threshold, corresponding to an angular deviation from horizontal of ∼14 degrees. Lever displacements were measured by optical sensors (Digi-key QRE1113-ND). Three cameras (Raspberry Pi Camera Module V2) recorded videos from each side of the box, and from the top (Extended Data Fig. 1a, Extended Data Fig. 5a).

#### Behavioral training

Water deprived rats received four 40-minute training sessions during their subjective night, spaced 2 hours apart. Starts of sessions were indicated by blinking house lights, a continuous 1kHz pure tone, and a few drops of water. At the end of each night, water was dispensed freely up to the daily minimum (5ml per 100g body weight).

Importantly, we wanted our paradigm to distinguish motor automaticity from habit formation^11,12^, two independent processes that can occur alongside each other and that may involve some of the same neural substrates^11,127,128^. Thus, we designed our task to ensure that rats achieve automaticity on a motor sequence without developing it into a habit. Since habits tend to form when the correlation between actions and outcomes is weak or variable^129–131^, if the reward is delayed^132^, or, further, if the reward is appetitive or addictive^133–135^, our paradigm directly linked behavioral variants to a water reward in a training process^131^ that resulted in highly overtrained behaviors expressed in goal-directed ways.

##### Stages of training

1. Rats were initially pre-trained to associate a visual cue with a lever press. On each trial, one of three LEDs above each of the three levers, chosen randomly, would light up to signify the ‘correct’ lever. Correct presses were rewarded while incorrect presses triggered a 1.2 second timeout, which was retriggered for every press in the time out period. To prevent rats from only selecting a single lever, we gradually decreased the probability of cueing a repeatedly pressed lever. All rats (n=18) learned to associate levers with cues in a median of 5284 trials. The criterion for learning was performing at >90% success rate for >100 trials.
2. After learning to associate visual cues with levers, rats were rewarded only when performing consecutive lever presses. Initially rewards were provided for every two successful consecutive presses, but after 500 rewards, reward was dispensed only after every three consecutive lever presses. Cued levers were constrained to not repeat, giving 12 different possible three-lever sequences. Rats quickly learned to press levers in a sequence, and no longer visited the reward port in between consecutive presses.
3. Following 1000 successful three-lever trials, rats were introduced to the block structure (Fig. 2a), which was modeled on a sequence task in primates^78^. In the block structure, the same three-lever sequence is cued on each trial until there are 6 successful performances, then a new sequence is randomly chosen. Cues are presented sequentially with no delay.
4. After ∼1-2 weeks of training on the block structure, working memory (WM) trials were introduced by withholding the 2^nd^ and 3^rd^ cued lever in the 4^th^-6^th^ trial of the block. Missed uncued levers were not required to be repeated until successful for a new sequence to be chosen.
5. One of the four nightly controlled sessions was changed to an automatic session. In this session, rats were required to perform only a single three-lever sequence, chosen randomly for each rat. This sequence was initially fully cued. Cues were then removed in reverse sequence, from last lever to first lever, every time the rat performed at >50% success rate over 30 trials. If success rate fell below 20% for 30 trials, or if rats failed to press the lever once in the entire session, a cue would be added back in. All rats were able to perform > 100 trials with at least two of the three cues withheld after 1024 ± 438 (mean ± SEM) trials or 17 ± 10 (mean ± SEM) days. After learning the sequence without cues, they would occasionally (5.18% ± 1.39% (SEM) of trials) hit the lower threshold prompting the addition of cues.

#### Behavior analysis

In total, 23 rats were trained on the three-lever task. 12 of 23 were used to characterize the behavior; nine were excluded because kinematics was not captured or analyzed early in training.

##### Definition of expert performance

Expert performance was determined when success rates and trial times stabilized to within 0.5σ of final performance values (based on the last 2000 trials). Metrics (success rates and trial time) were smoothed with a moving average of 400 trials. Furthermore, we required automatic performance to reach >72% success rates to be considered ‘expert’, following previous studies^55,85,87,125^.

##### Calculation of performance metrics

###### Success rate

Success rate was defined as the number of rewarded trials divided by the total number of attempted trials.

###### Trial time

Defined as the interval between the first and third lever press. This only includes successful sequences, as incorrect sequences may not include three full lever presses.

###### Error variability

Defined as the Shannon entropy (in bits) of the probability of each sequence occurring for a given target sequence. Low probability sequences (p<0.001) are discarded. If mistakes are systematic, the probability distributions will be skewed towards particular erroneous sequences, and the entropy will be low. If mistakes are made randomly, the distribution will look more uniform, and the entropy will be high. For controlled sequences, the error calculation was done on the sequence chosen for the OT mode.

###### Error modes

Motor errors were defined as failures to touch or fully depress the ‘correct’ lever in what would otherwise have been a correct sequence (Supplementary Movie 2). I.e. the rat oriented to the ‘correct’ lever and swiped at it but failed to depress it beyond the threshold for detection. Sequence errors, on the other hand, involved orienting to and pressing the wrong lever. For each rat and session type (controlled and automatic), ∼100 videos of error trials were manually inspected and labeled as either a sequence or motor error. To analyze behavior following different error mode types (Fig. 3f), we used a heuristic to automatically estimate and classify failures as motor or sequence errors. This allowed us to analyze >100 trials. In this analysis, motor errors were classified as any error sequence that resembled an omitted lever (e.g., for the target sequence LRC, RC and LC are considered motor errors), while sequence errors are any other type of mistake. This heuristic generally overestimates motor errors (29.65% ± 5.9% of errors for CUE trials, 16.95% ± 5.48% of errors for WM trials, 13.16% ± 5.32% of errors for OT trials, data is mean ± s.e.m.) and underestimates sequence error. Trial-dependent accuracies and trial times, conditioned on the type of trial that came before (hit, motor error, sequence error), were then calculated using this heuristic (Fig. 3f). For trial times, we only considered sequences that had the same overall movement length as the overtrained sequence (since L→R is further to travel than L→C) as a control.

###### Average forelimb speed

Raw trajectories (position traces) of the active forelimb were smoothed and up-sampled (from 40hz to 120hz) using cubic smoothing spline (csaps in Matlab, smoothing parameter of 0.1). Instantaneous velocities for the horizontal (x) and vertical (y) positions were calculated, then converted to instantaneous speed. This value was averaged from 0.1 second before the first lever press to 0.1 second after the last one. Since velocity was calculated from a side-view camera, and animals moved towards and away from the camera to press different levers, we left velocity measures in pixels/s.

###### Movement smoothness

To quantify the smoothness of the movement, or its continuity and non-intermittency^71^, we measured each trajectories spectral arc length, a dimensionless metric that measures the arc length of the Fourier magnitude spectrum within an adaptive frequency range^103^. This metric quantifies smoothness independent of amplitude and duration, and is less sensitive to noise than another popular smoothness metric, the log-dimensionless jerk^71,136^. Values are scaled between the maximum and minimum average spectral arc length recorded across rats in Figures 2i, 6i, and 7i. Values closer to 1 are more smooth.

##### Trial selection for behavioral analysis

In Fig. 2, early performance is taken from the first 1000 trials, and late performance is measured from the last 1000 trials. For Fig. 3, metrics are calculated from all trials after expert performance was reached. For Fig. 6 and 7, pre- and postlesion accuracies are calculated from the week before and after lesion. Pre- and postlesion trial times, trial speeds, and kinematics are taken from the last 1000 trials before and after lesion. Finally, it is important to note that since automatic sessions are introduced after the controlled sessions, early performance on automatic (OT) trials benefits from prior controlled practice (Fig. 2).

##### Kinematic tracking

To track the movements of the rat’s active forelimb and head during our task, we utilized recent machine learning approaches to detect keypoints from individual video frames. Videos of animals performing the task were acquired at 40 Hz (90 Hz for DLS recording cohort) from cameras pointing at the lever from the two sides to obtain both forelimb trajectories, and one camera pointing down from the top to obtain the rat’s horizontal position. For video-based tracking, we trained ResNet-50 networks pretrained on ImageNet, using DeeperCut (https://github.com/eldar/pose-tensorflow)^82^. To refine the tracking for our rats, we randomly selected about ∼200 frames per view and trained the network using manually labeled position of the hand and nose. The network was then used to predict the position of body parts across all trials on a frame-by-frame basis, using GPUs in Harvard Research Computing cluster. Tracking accuracy was qualitatively validated post-hoc by visual inspection of 5 trials across 3 different sessions. Frames with poor tracking (< 0.95 score from the model), due to occlusion of the forelimbs, were removed and trajectories in those frames were linearly interpolated. If > 5 consecutive frames were removed, the trial was discarded for tracking purposes. Additionally, any trial with >5% of poorly tracked frames was removed from the analysis. The full trial trajectory was then smoothed using a Gaussian filter in matlab, with a σ of 0.6 frames.

To track the movements of the rat’s active forelimb and nose in 3-dimensions, we first calibrated our multiple camera views (left side, right side, top side, see Extended Data Fig. 5a) to a set of manually labeled features in our box observable from both views in order to calculate camera extrinsics and world coordinates, drawing from camera calibration functions in the Computer Vision Toolbox (e.g. estimateWorldCameraPose, estimateCameraParameters, cameraPoseToExtrinsics). We could then use the calibrated cameras to triangulate our 2D estimated points into 3D^137^. For the forelimb, we triangulated from 2D points tracked on the left and right cameras (Extended Data Fig. 5b). The nose used either the left and top, or right and top cameras. 3D world coordinates are in mm, relative to one of the manually labeled feature in the box.

##### Kinematic analyses

To quantitatively compare kinematic similarity, we computed pairwise trial-to-trial correlations. Since trial times varied, movement trajectories were time warped to a common template. Specifically, trajectories from each trial were interpolated so that the time between the 1^st^ and 2^nd^ lever, and the time between the 2^nd^ and 3^rd^ lever, matched the median inter-lever intervals. For sub-movement correlations, trajectories are warped to the median inter-lever interval time. Though we tracked both forelimbs, analyses were performed only on the active forelimb used to press the lever. Rats used a single forelimb to perform lever presses (n=12/23 right, n=11/23 left).

#### Electrophysiological recordings

Microdrive construction, surgical and recording procedures were performed as previously described^138^. After expert performance was reached on the sensory guided, memory guided, and automatic tasks, a microdrive containing arrays of 16 tetrodes was implanted into the DLS (n=4 rats) contralaterally to the active forelimbs as previously described^24^ (Extended Data Fig. 2a). Neural and behavioral data was recorded continuously for 95+-31 days. The drive was occasionally advanced by ∼200 µm, 0-4 times over the course of the experiment. At the end of the experiment, an electroylytic lesion was done to mark the electrode site. This was done by passing a 30 µA anodal current for 15s through the electrode tips. For implant coordinates, according to Paxinos^139^, see Table 1.

**Table 1:** Surgical coordinates and injection volumes.

|  | DLS lesion |  |  |  | DLS implant | DMS lesion |  |  |  |
| --- | --- | --- | --- | --- | --- | --- | --- | --- | --- |
| AP | +0.7 | +0.7 | -0.3 | -0.3 | 0.5 | +1.2 | +1.2 | +0.2 | +0.2 |
| ML | ±3.9 | ±3.9 | ±4.1 | ±4.1 | ±4.0 | +2.2 | +2.2 | +2.0 | +2.0 |
| DV | -4.5 | -3.5 | -4.8 | -4.0 | 4.0 | -4.4 | -3.4 | -4.4 | -3.4 |
| nL | 25*4.5 | 19*4.5 | 20*4.5 | 16*4.5 |  | 25*4.5 | 25*4.5 | 25*4.5 | 25*4.5 |

#### Lesion surgeries

After reaching expert performance, bilateral striatal lesions were performed (n=7 DLS, n=6 DMS) as previously described^24^. For injection coordinates, see Supplementary Table 1. In brief, quinolinic acid (0.09 M in PBS (pH 7.3), Sigma-Aldrich) was injected in 4.5nl increments, via a thin glass pipette connected to a microinjector (Nanoject III, Drummond). Lesions were performed in two stages, starting with the side contralateral to the primary forelimb (the forelimb that presses the first lever). Animals recovered for 7 days before being reintroduced to training.

#### Histology

At the end of the experiment, animals were euthanized (100 mg/kg ketamine and 10 mg/kg xylazine), transcardially perfused with either 4% paraformaldehyde (for nissl staining to confirm lesion size and location), or 2% paraformaldehyde (PFA) and 2.5% glutaraldehyde (GA) (for osmium staining, to confirm location of electrode implant) in 1x PBS. For electrode implants, brains were then stained with osmium (as described in^140^) and embedded in epoxy resin for micro-CT scanning. Micro-CT scans (X-Tek HMS ST 225, Nikon Metrology Ltd.) were taken at 130 kV, 135 uA with 0.1 mm copper filter and a molybdenum source. 3D volume stacks were reconstructed (VG studio max), and brains were aligned along the coronal, medial, and sagittal plane using Fiji. Location of the electrolytic lesion could be calculated relative to anatomical landmarks (i.e., corpus callosum split at AP = 1.65mm from bregma, anterior commissure split at 0 mm bregma). For lesioned animals, brains were sectioned into 80-µm slices using a Vibratome (Leica), then mounted and stained with Cresyl-Violet. Images of whole brain slices were acquired at x10 magnification with either a either a VS210 whole slide scanner (Olympus) or an Axioscan slide scanner (Zeiss). To quantify the extent and location of striatal lesions, we analyzed coronal sections spanning the anterior posterior extent of the striatum from 4 calibration animals and 2 experimental animals (DLS) (7 hemispheres injected in total) or from 4 experimental animals (DMS). Boundaries were manually marked based on differences in cell morphology and density^141,142^. The extent of the striatum was determined based on the Paxinos^139^, using anatomical landmarks (external capsule, ventricle) and cell morphology and density.

#### Neural analysis

##### Spike sorting

Raw neural data was collected continuously over the course of the experiment (mean and s.e.m. are 95 ± 31 days, n=4 fully trained rats). Spiking activity from populations of single units was sorted using our custom-designed spike-sorting algorithm, Fast Automated Spike Tracker (FAST)^138^. A custom Matlab GUI (https://github.com/Olveczky-Lab/FAST-ChainViewer) was used to manually isolate and track single-units over long timescales. On average, we isolated 20.5 ± 13.3 units simultaneously in the striatum within each session. We were able to track units for an average of 4.3 ± 1.2 days. Assessing the quality of sorted single unit was done as previously described^138^. In brief, we evaluated unit quality by computing the isolation distance^143^, the L-ratio^144^, the fraction of inter-spike interval violations^145,146^, discarding units that did not meet our criteria (Isolation distance >=25, L-ratio <=0.3, and ISI violations below 2m <= 1%).

##### Unit type identification

Units were identified as putative MSNs or FSIs based on their peak width (full width at half maximum) and time interval between spike peak and valley^147,148^. Units with peak width >150 µs, peak-valley interval >500 µs were classified as SPNs, while units with peak width ≤150 µs, peak-valley interval ≤500 µs were classified as FSIs.

##### Criteria for unit selection

We selected a subset of the total population of recorded units for our neural analyses. To be included, we required that a neuron fired at least 1 spike, on at least 25% of all trials and was recorded over >5 rewarded trials in each execution mode (CUE, WM, OT) for Fig. 4, or >5 rewarded trials in each of the 12 different sequences. From a total of 2468 recorded and well-isolated units, this criterion reduced the units available for analysis to 579 units for the comparison across execution modes, and to 340 units for comparisons across sequences in the controlled sessions.

##### Neural metrics

###### Trial averaged, z-scored activity

First, instantaneous firing rates were calculated for each trial by convolving binned (10ms bins) spike counts with a Gaussian kernel (σ=25ms). To account for differences in trial times, firing rates were then local linearly warped to the median press times. Warping was done only after calculating firing rates so as to not artificially increase or decrease the firing rate. Firing rates before the first and after the last lever press were not warped. After this alignment step, each trial was z-scored, and then averaged over for each unit and each execution mode.

###### Average firing rates

Firing rates on individual trials were calculated from -0.2 seconds before the first lever press, to 0.2 seconds after the third lever press. This value was averaged over every trial for a given execution mode.

###### Average activity

To determine if there was elevated population activity at privileged timepoints in the sequence in the task (e.g. at the boundaries of discrete motor elements), we averaged over the time varying z-scored activity for each unit recorded in a rat. This average trace was compared to a distribution of average z-scored activity, sampled from random times in the behavior (2 seconds before and after the first and last lever press, n=1 × 10^4^ permutations).

###### Correlation across execution modes

For each unit, correlation coefficients were computed between the time-varying vector of trial-averaged activity across execution modes.

###### Correlating neural activity associated with behavioral elements across sequences

We computed the correlations between neural population vectors of combined orienting and lever pressing movements across the 12 different sequences. Population vectors were calculated by averaging the activity of each neuron over orienting and lever press movements. Orienting movements were defined as those that occurred 0.1 seconds after a prior lever press until 0.1 seconds before the next press. Lever press movements were defined as occurring +/-0.1 seconds around the lever deflection. We excluded the first lever press for this analysis, as it was not preceded by an orienting movement.

###### Principal components

Principal component analysis was performed on the matrices of population activity (neurons vs. time) concatenated across either the three execution modes (CUE, WM, OT) or the twelve sequences along the time dimension. Execution modes or sequences were then disjoined to generate the plots in Fig. 4e and Fig. 5d.

##### Neural decoding analysis

We used a feedforward neural network with two hidden layers to predict the time-varying, 2D velocity components of the active forelimb (side camera) and the nose (top camera) from the spiking activity of ensembles of DLS neurons. Spiking activity was binned in 25ms bins. We used 75ms of coincident spiking activity as the input to the model. Other model parameters were the same as in previous work^24^. We additionally challenged our network to predict the 3D velocity components of the active forelimb and nose, from 3D world coordinates triangulated from calibrated cameras (described above) (Extended Data Fig 5d-f).

We trained our models on blocks of >50 trials in which there were at least 12 simultaneously recorded units that had an average firing rate >0.25hz during the trials. In each block of trials, we fit decoding models using the activity of up to n=20 randomly sampled ensembles of 12 striatal units. We quantified model performance using two-fold cross validation by computing the pseudo-R2: 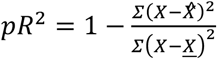 Decoding performance (pseudo-R2) was measured in each ensemble, then averaged across all 20 ensembles, and then averaged across all blocks of trials for each rat. For subset model, the training dataset was generated from only 6 of the 12 sequences, chosen at random for each of the n=20 ensembles. The test dataset was then generated from the remaining 6 sequences.

### Neural network model

We simulated an artificial neural network consisting of two populations, one corresponding to DLS and another to other downstream motor circuits. The DLS network contained no recurrent connections while the downstream network contained all-to-all recurrent weights. The two populations were bidirectionally connected with all-to-all feedforward and feedback weights. The downstream motor network directly controls movement via a set of feedforward weights, and also receives an additional source of input representing cue signals via a set of feedforward weights. Each network consists of 500 units with a rectified linear (ReLU) activation function. Network weights were initialized with the Kaiming Uniform initialization^149^. Gaussian noise of standard deviation 0.1 was added to the inputs to each neuron in both networks at every timestep in all simulations.

We modeled a simplified version of the experimental task in which the output of the network controls the velocity of a “forelimb” (represented simply as a point) and is tasked with moving it into a set of three circular target zones in a prescribed sequential order, as in the piano task. The target zones were positioned as shown in Fig. 8b. On each trial, the loss function measuring the performance of the network was defined as the sum of the squared distance between the forelimb position and the center of the current target. The identity of the current target changes to the next in the sequence once it is reached. On cued trials the target lever changed, and on the first step, cue input was provided in the to the downstream network in the form of a vector indicating the position of the cue relative to the forelimb. The cue input was transient, lasting only one timestep for each cue. If the sequence was not performed successfully within T=40 timesteps of the simulation, the trial was halted and considered a failure.

The network, excluding the DLS input and output weights, was pre-trained on the cued task for 100,000 iterations (well past the point where asymptotic performance was reached). The DLS input weights were then trained on randomly interleaved cued and automatic task trials (50% probability of each, with all 12 possible target trajectories equally likely on cued trials), again for 100,000 iterations. The target sequence on automatic trials was always the same (right, center, left). All network training used backpropagation and the Adam optimizer with learning rate set to 1e-4. Training was conducted using PyTorch.

In Extended Data Fig. 8a-g, we modified the network architecture by replacing the NxN DLS output weights with a chain of Nx1 and 1xN weights, corresponding to a rank-1 projection. In Extended Data Fig. 8h-m, we modulated the gain of DLS activity, suppressing it by a factor of 0.1 at all time steps except the first and those when the target lever changed. In Extended Data Fig. 8n-s, we omitted the pretraining stage and instead trained the entire model on all task modes for 200,000 iterations.

## Extended Data

**Extended Data Figure 1:**
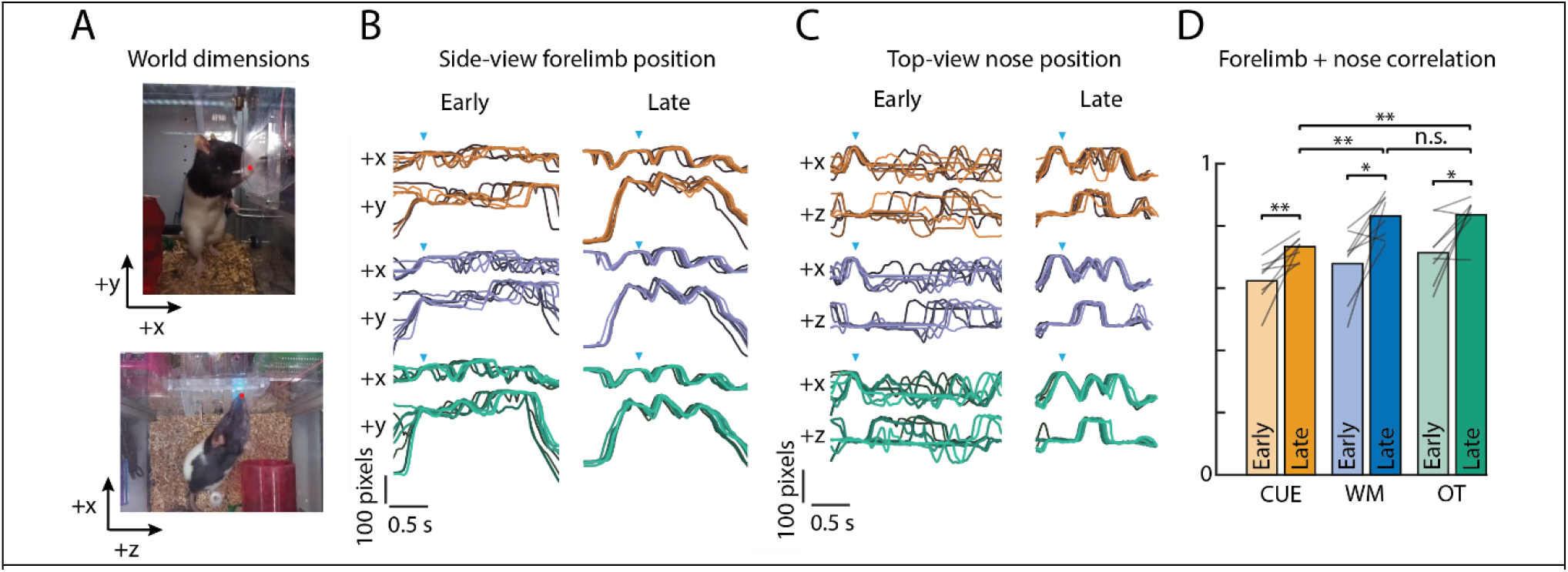
Rat’s movements increase stereotypy along other axes and joints. **A**. View of ‘piano task’ from a side (right) and top camera. Axes are defined as +x – towards lever, +y – towards top of box, and +z – towards right lever along the piano. **B**. Replotted from Fig 1e is 8 example forelimb trajectories in the x and y dimension for each execution mode from early and late in learning. Orange – CUE, blue – WM, green – OT. **C**. Same as B., but for the nose position in the x and z dimension. **D**. The average, trial-to-trial correlation of forelimb (in x and y dimensions) and nose (in x and z dimensions) trajectories increases with training. Bars represent grand averages over rats, and lines are averages within individual rats.

**Extended Data Figure 2:**
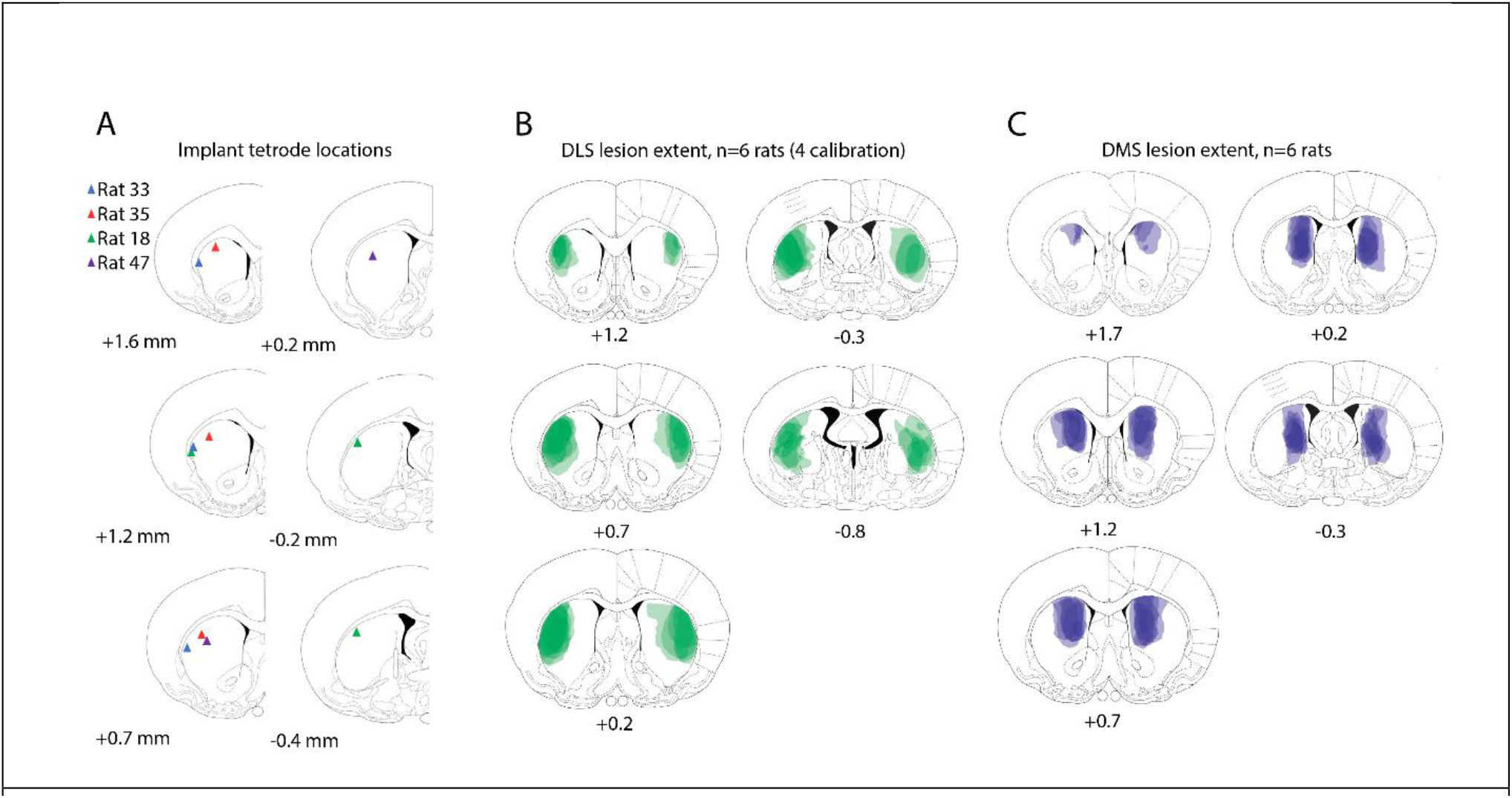
Histology of DLS implants, DLS lesions, and DMS lesions. **A**. Location of recording electrode implantation sites in DLS marked with a colored arrowhead for each of the 4 rats. For some individuals multiple sites are marked, due to individual tetrode bundles spreading during implantation. Coronal slices are labeled from distance relative to bregma. **B**. The extents of DLS lesions from 6 rats (11 hemispheres) are marked for the DLS lesion along the anterior-posterior axis of the striatum, and shaded in green. Lesion extent was calibrated to target the motor cortex-recipient region of dorsolateral striatum, as determined from virally-mediated fluorescent labeling in^24^. **C**. Same as B, but 12 across 6 rats hemispheres are labeled for DMS lesions. Targeting is based on the prefrontal cortex recipient region of dorsomedial striatum, also from work in^24^.

**Extended Data Figure 3:**
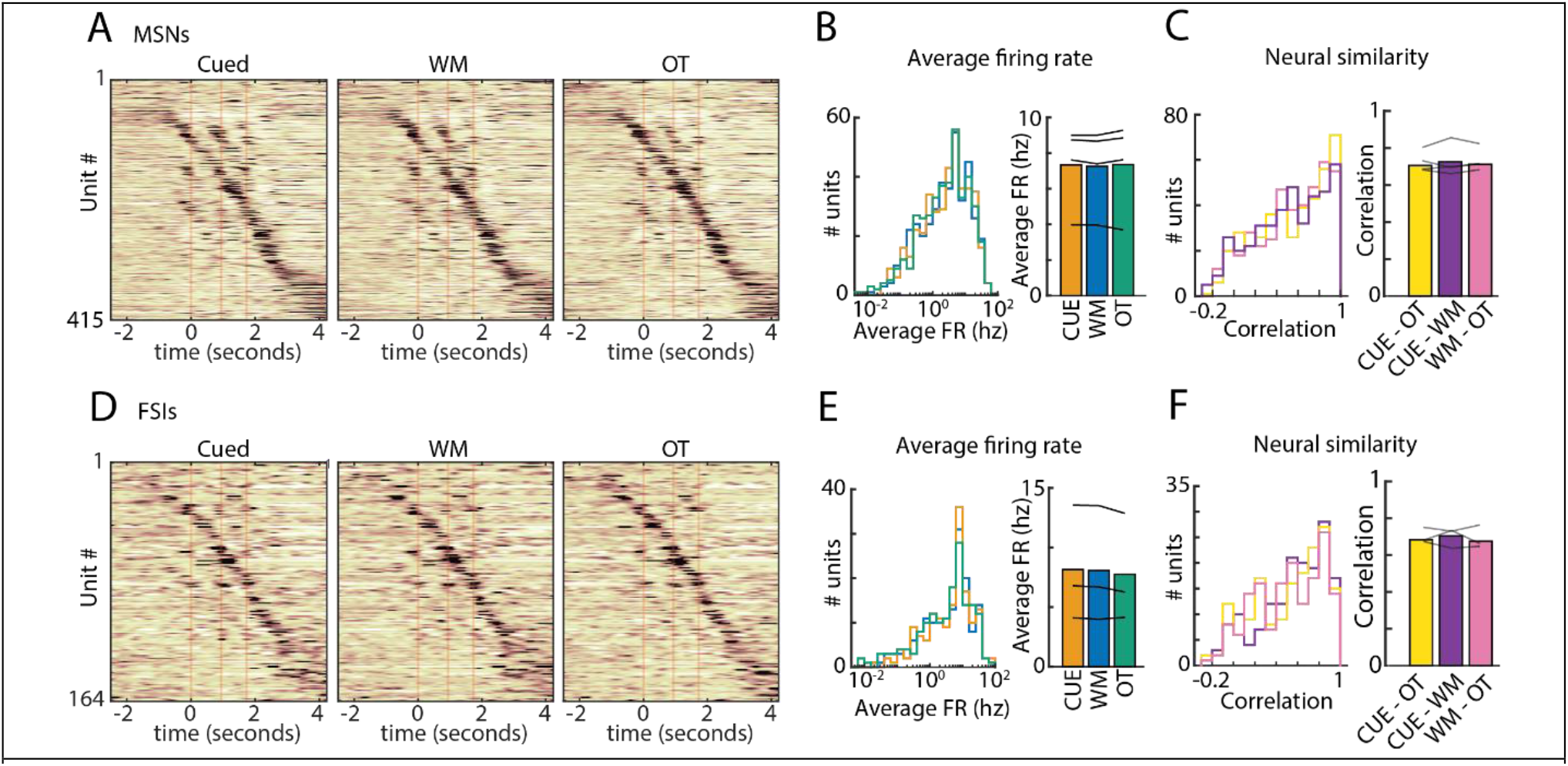
MSNs and FSIs represent OT, CUE, and WM sequences similarly. **A**. Z-scored average activity of 415 putative medium spiny neurons (MSNs) recorded in the DLS for the same sequence during the OT, CUE, and WM execution mode (from n=4 rats). The trials were linearly time-warped to each lever press (red vertical lines). Units were sorted by the time of their peak activity. The sorting index was calculated from half the available trials for each unit, taken from the OT mode, and then applied to the remaining trials and modes. **B**. (Left) Histogram of the average firing rate of putative MSNs during the trial period for each execution mode. (Right) Average firing rates across all rats are not significantly different (p>0.05, two-tailed t-test). Lines represent individual rats. **C**. (Left) Histogram of correlation coefficients of trial-averaged neural activity across the execution modes (CUE x WM - purple, WM x OT - pink, CUE x OT - yellow). (Right) Average correlation coefficient across all units, for each rat (n=4). Average correlations are not significantly different across each mode comparison (p>0.05, two-tailed t-test). **D-F**. Same as A-C, but for putative FSIs. Note n=3 only, as one rat had no putative FSIs recorded that met our criteria (see Methods).

**Extended Data Figure 4:**
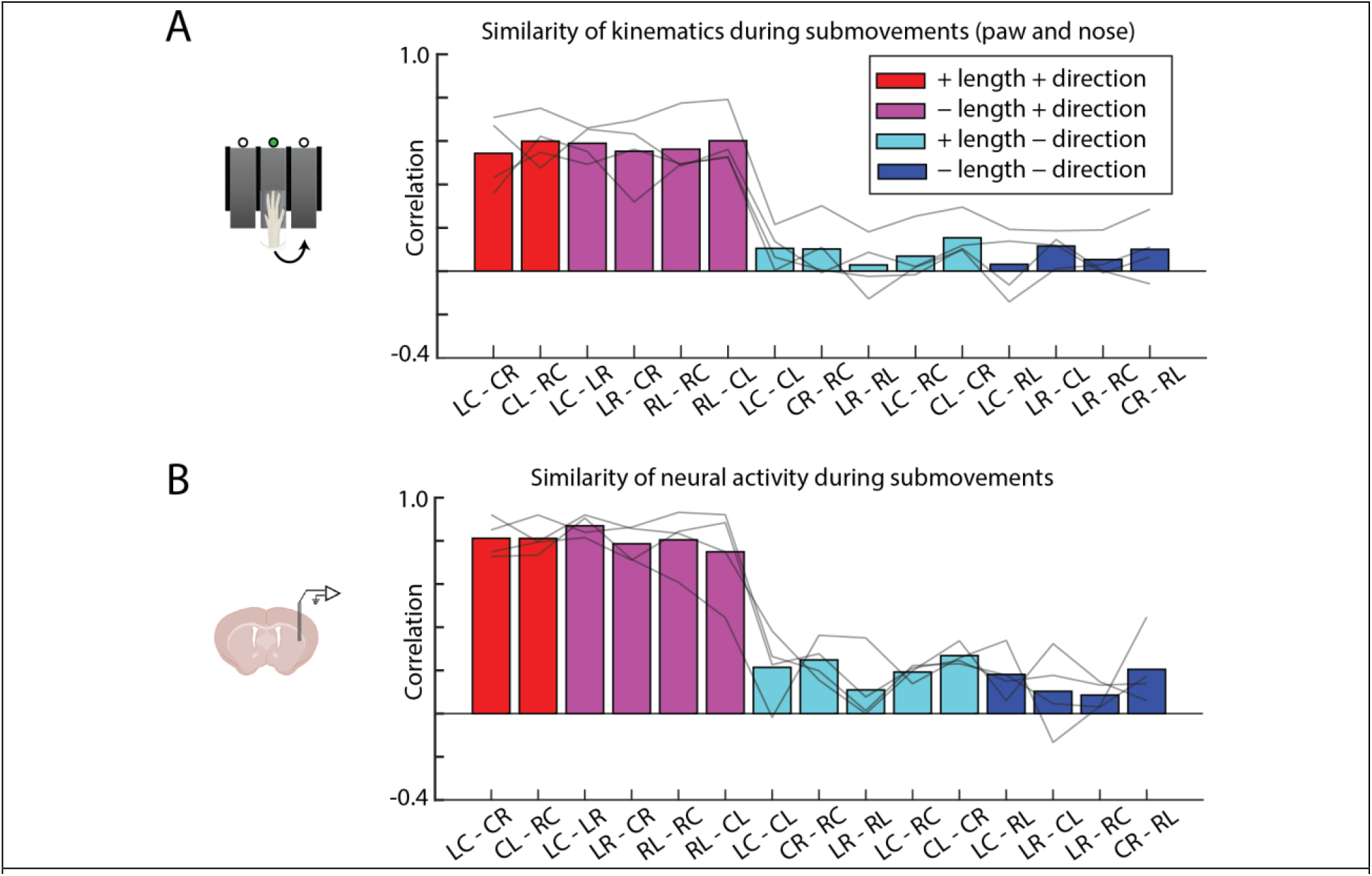
Neural and kinematic similarity for all orientation movements. **A**. Comparing kinematic similarity across different orientation movements. Plotted is the average trial-to-trial correlation between kinematic traces of the forelimb (side view, x and y) and nose (top view, x and z) from different orientation movements (e.g., L->C and C->R). Orientation movements are cropped 0.2 seconds after and before the lever presses. Bars are averages across rats, and lines represent averages in individual rats (n=4). Colors denote whether orientation movements match in length (i.e. short vs. long) or orientation direction (i.e. left-vs. right-wards). B. Comparing neural similarity across different orientation movements. Population activity is averaged during the orientation movement (defined as 0.2 seconds after and before the presses) for each different orientation movement, and correlation coefficients are computed between population vectors.

**Extended Data Figure 5:**
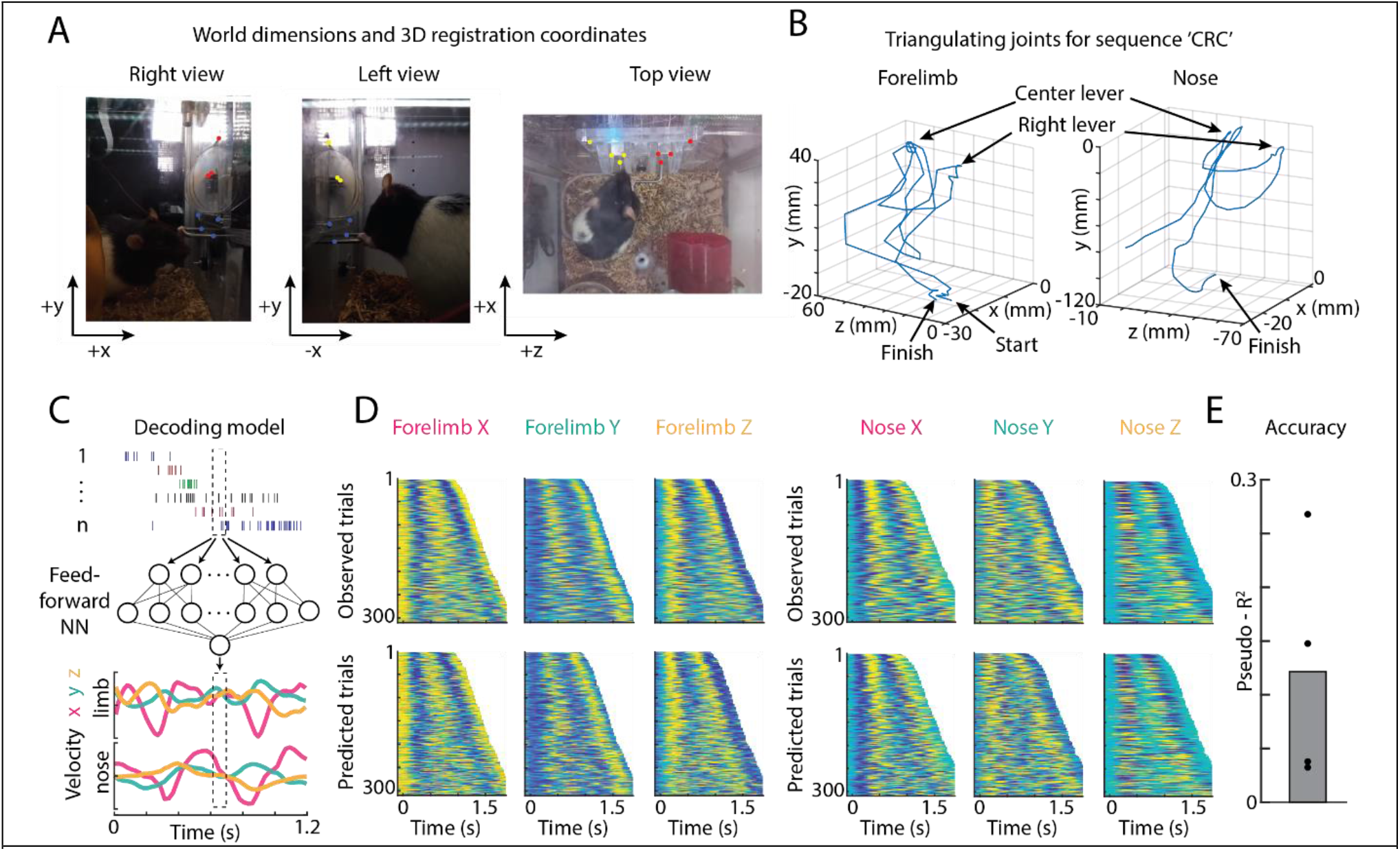
DLS encodes 3d nose and forelimb kinematic trajectories. **A**. Views from our three cameras (right, left, and top) are shown, along with a set of static features in the box that were used to calibrate multiple views to the world for triangulation^137^. To triangulate the forelimb, the left and right view were calibrated using the blue points. To triangulate the nose, the top and either left or right view were calibrated using the yellow or red points. Some points are shared across calibrations. **B**. An example trajectory of the forelimb (left) and nose (right) plotted in 3 dimensions, during performance of the sequence C->R->C. Forelimb coordinates are relative to the top-left blue point in A, and nose coordinates are relative to the top-left yellow point in A. **C-E**. Decoding analysis, performed the same as in Fig. 4f-h. **C**. Schematic of the decoding analysis. A feed-forward neural network is trained to predict the velocity components (x, y, and z) of the nose and forelimb in 3 dimensions. **D**. (Top) Heatmap of normalized forelimb (left) and nose (right) velocities in each dimension, observed in an example controlled session. (Bottom) Heatmap of the predicted forelimb and nose velocities output by our model. **E**. Decoding performance, measured in pseudo-R^2^ of the model on a held-out set of test trials (see Methods). Dots indicate model performance on individual rats, and bar is average over rats (n=4).

**Extended Data Figure 6:**
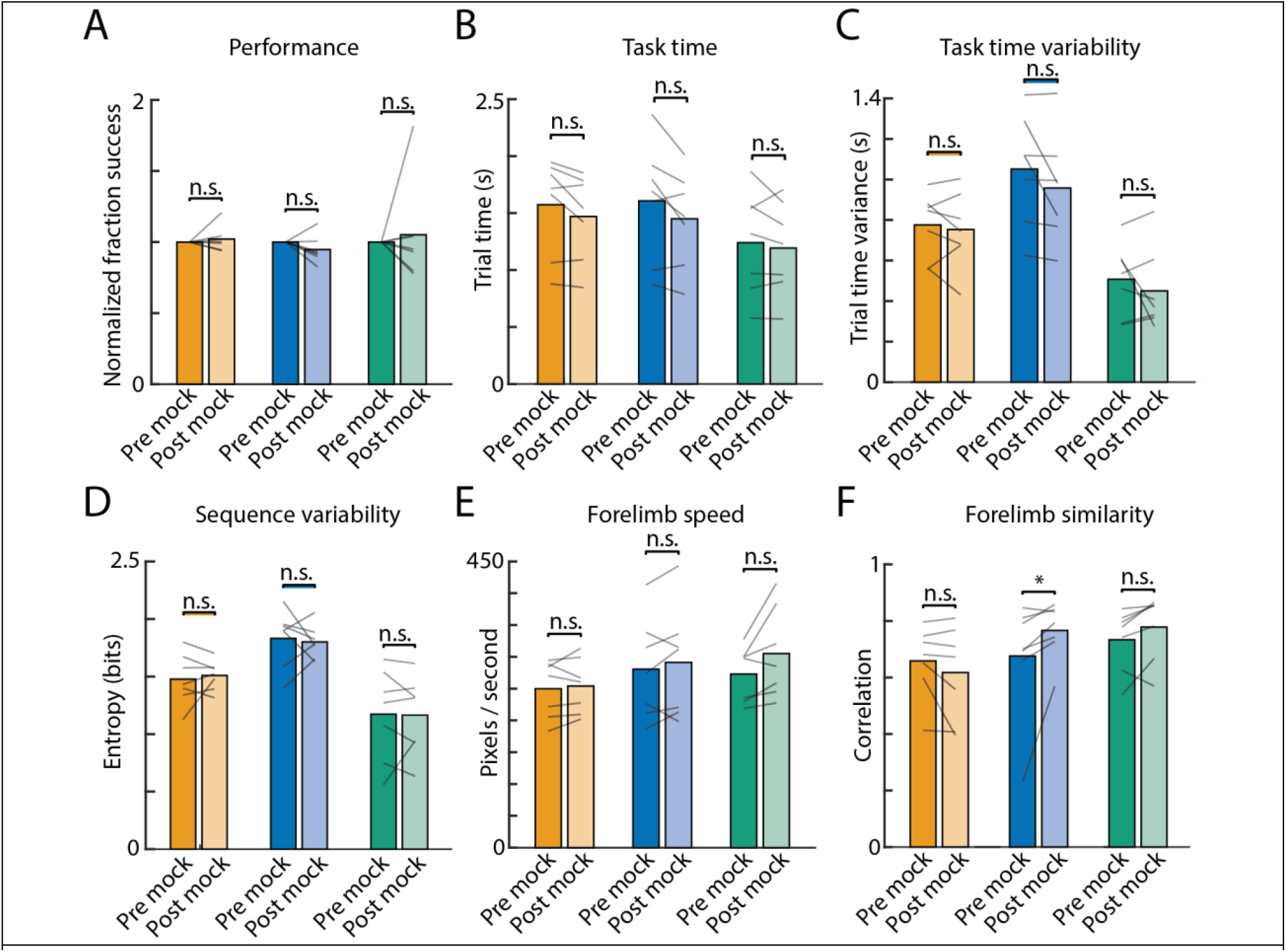
Performance on 3-lever task is unaffected by a 7 day mock break. Performance metrics before and after the mock break, in expert animals. Gray lines represent individual rats, bars are averages across rats. **A**. Normalized success rate, **B**. Trial time, **C**. Variance in the trial time, **D**. Entropy, or randomness, of errors, **E**. Average speed during the trial, **F**. Average trial-to-trial correlation. *P<0.05 Wilcoxon signed rank test.

**Extended Data Figure 7:**
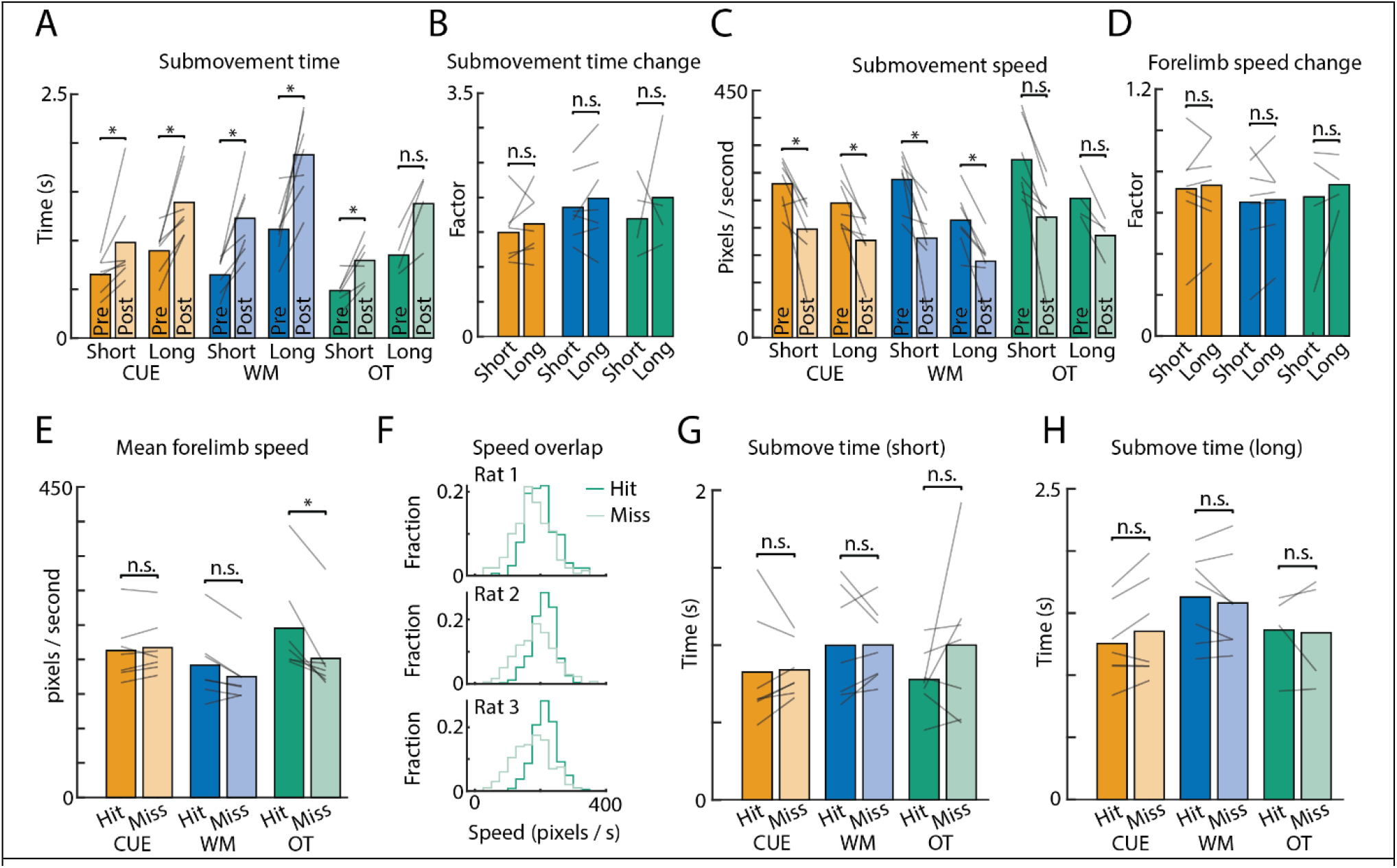
DLS lesion affects different submovements similarly. **A-D**. DLS lesions affect short (e.g., L->C) and long (e.g., L->R) orientation movements similarly. **A**. Plotted is the average inter-lever interval, split by short and long orientation movements, before (darker shade) and after (lighter shade) the lesion, for each execution mode. Note that only 4 of 7 rats had long orientation movements in their prescribed OT sequence. In all plots, lines represent averages within individual rats, and bars are grand averages over all rats. **B**. The factor increase in trial time (post lesion time / pre lesion time) is similar for short and long movements. **C-D**. Similar to A-B, but for the average forelimb speed during the orientation movements (submovements). **E-H**. Successful performance postlesion is not completely dependent on speedy execution of the motor sequence. **E**. Average forelimb speed during all successful (Hit) and unsuccessful (Miss) trials, following the DLS lesion. **F**. Distributions of forelimb speeds (x-axis) for successful (darker shade) and unsuccessful (lighter shade) trials for three example rats. **G-H**. Average inter-lever interval (submovement time) for actions performed in successful (Hit) and unsuccessful (Miss) trials. **G**. For short (e.g., L->C) submovements, and **H**. long (e.g., L->R) submovements. Again, only 4 of 7 rats had a long orientation movement in the OT sequence. *p<0.05, Wilcoxon sign rank test.

**Extended Data Figure 8:**
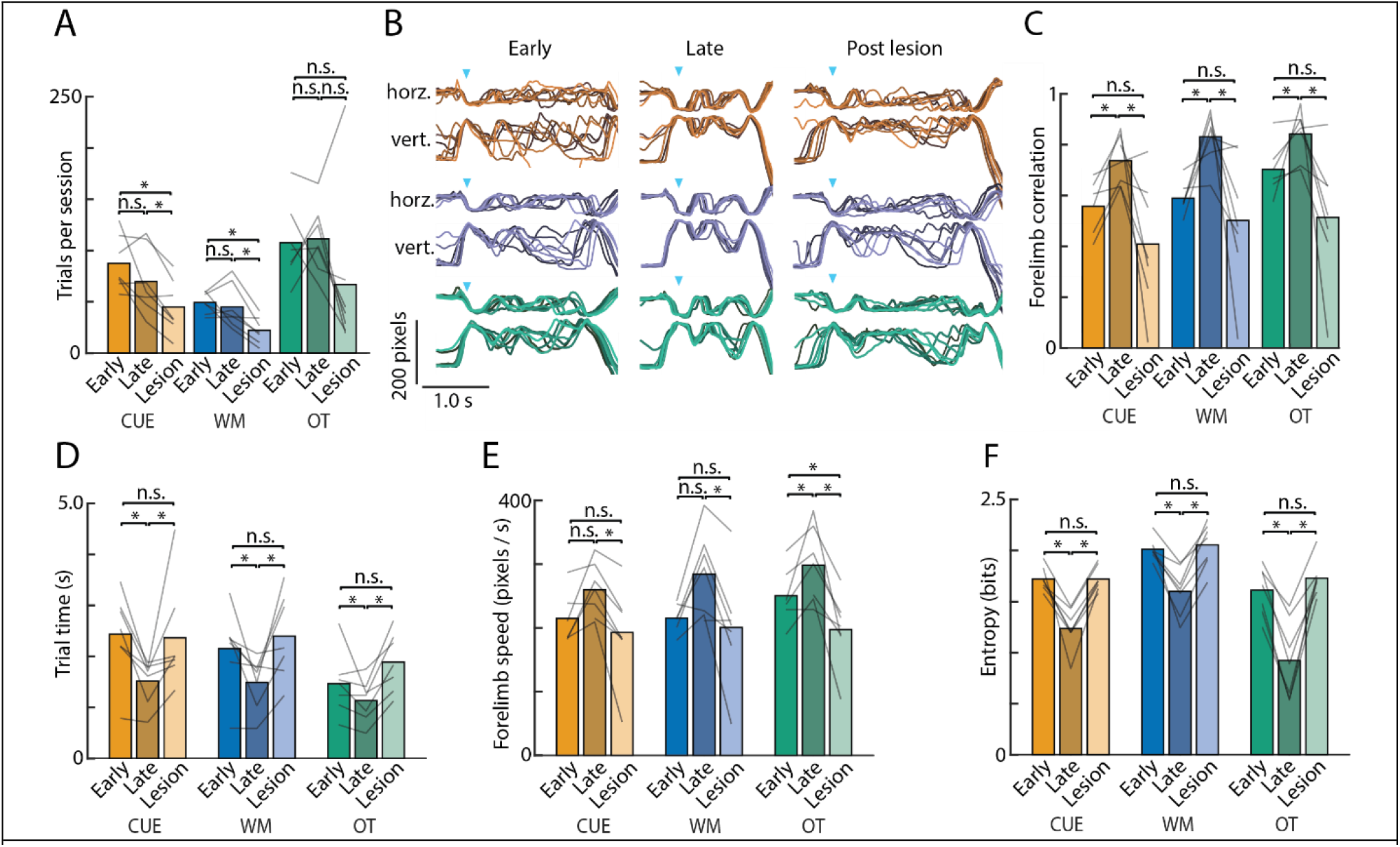
Post-lesion kinematics are more similar to early in learning. **A**. Trials presses per session for both CUE, WM, and OT sequences decrease on average following the lesion. Average number of trials per session was not significantly different between controlled (CUE, WM) and automatic (OT) session types before (P=0.9375) or after (P=0.8125) the lesion. **B**. Forelimb kinematics from 8 example trials sampled early in learning, late in learning, and following the bilateral DLS lesion (also see Fig 1e, Fig 5e, and see Methods for timing). **C**. Average trial-to-trial correlation for forelimb trajectories of the active paw (both horizontal (x) and vertical (y)) from early in training, compared to late (pre-lesion), and post-lesion, for all execution modes (red=CUE, blue=WM, green=OT). Gray lines are average within rats (n=7 late and lesion, n=6 early, 1 rat was not recorded early in learning) and bars represent average across rats. **D**. Trial time from 1st to 3rd lever press early, late (or pre-lesion), and post-lesion. **E**. Variance in the trial time. **F**. Average forelimb speed during the trial. *P<0.05, Wilcoxon sign-rank test.

**Extended Data Figure 9:**
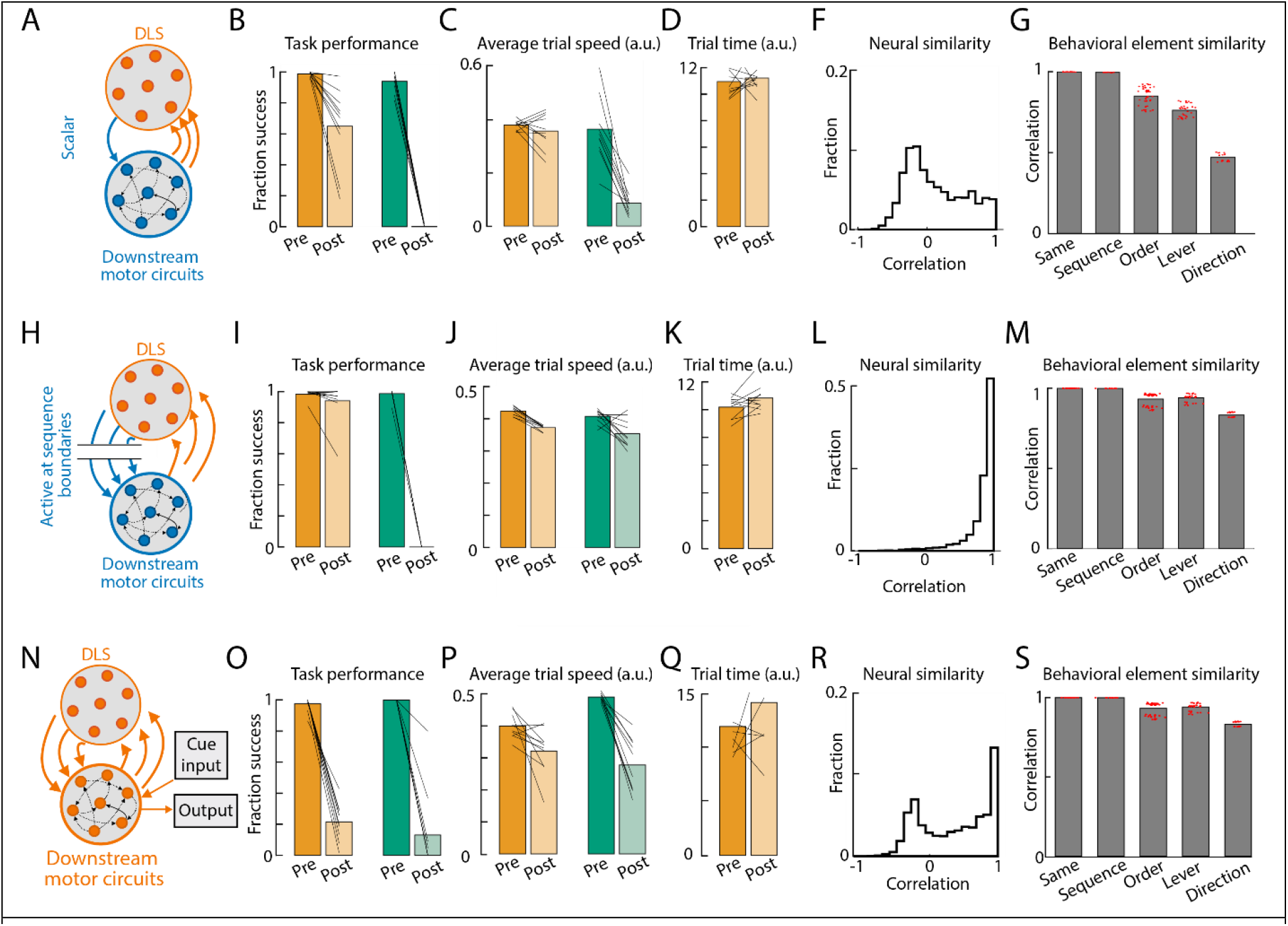
Alternative network models fail to reproduce experimental results. **A-G:** A neural network model with scalar DLS outputs fails to learn mode-invariant DLS activity **A**. Schematic illustrating architecture of a model variant in which DLS outputs to downstream motor circuits are constrained to be scalar-valued. **B-G:** Replication of analyses in Figure 8d,f-i, for this model variant. The neural representations are much less similar across modes than in the original model (panels E and F here versus Fig. 8d, f). **H-M:** A neural network model with action selection signals fails to learn strong kinematic representations. **H**. Schematic illustrating architecture of a model variant in which DLS outputs to downstream motor circuits are suppressed except at trial initiation and transitions between lever presses. **I-M**: Replication of analyses in Figure 8d,f-i, for this model variant. The neural representations show much less egocentricity than in the original model (panel M here vs. Fig. 8f). **N-S:** A neural network model without pre-trained circuits is not robust to DLS lesions in the controlled mode. **N**. Schematic illustrating architecture of a model variant in which the entire model is trained on the cued and automatic tasks from scratch, rather than using the strategy of pretraining downstream motor circuits on cued trials first. **O-S:** Replication of analyses in Fig. 8d,f-i, for this model variant. The resilience of controlled mode performance seen in the original model is lost (panel O here versus Fig. 8g).

## Supplementary Info

### Supplementary Video 1

Three trials of the same motor sequence performed in the CUE, WM, and OT execution mode. Videos are shown first from a top camera, then from a side camera. In the side videos, the kinematics of the active forelimb is tracked and plotted.

### Supplementary Video 2

Trials from the OT mode from an example rat are shown to demonstrate the types of errors we observe. Example trials include five consecutive successful trials, two motor errors that follow a successful trial, and finally a run of sequence errors following motor errors.

